# Deep learning, data ramping, and uncertainty estimation for detecting artifacts in large, imbalanced databases of MRI images

**DOI:** 10.1101/2023.06.20.545422

**Authors:** Ricardo Pizarro, Haz-Edine Assemlal, Sethu K. Boopathy Jegathambal, Thomas Jubault, Samson Antel, Douglas Arnold, Amir Shmuel

**Affiliations:** McConnell Brain Imaging Centre, Montreal Neurological Institute, McGill University, Montreal, QC, Canada; Department of Neurology and Neurosurgery, McGill University, Montreal, QC, Canada; Department of Biomedical Engineering, McGill University, Montreal, QC, Canada; Department of Physiology, McGill University, Montreal, QC, Canada; NeuroRx Research, Montreal, QC, Canada

**Keywords:** Deep transfer learning, Data ramping, Uncertainty metrics, Large neuroimaging database, Imbalanced datasets, Quality control

## Abstract

Magnetic resonance imaging (MRI) is increasingly being used to delineate morphological changes underlying neurological disorders. Successfully detecting these changes depends on the MRI data quality. Unfortunately, image artifacts frequently compromise the MRI utility, making it critical to screen the data. Currently, quality assessment requires visual inspection, a time-consuming process that suffers from inter-rater variability. Automated methods to detect MRI artifacts could improve the efficiency of the process. Such automated methods have achieved high accuracy using small datasets, with balanced proportions of MRI data with and without artifacts. With the current trend towards big data in neuroimaging, there is a need for automated methods that achieve accurate detection in large and imbalanced datasets. Deep learning (DL) is the ideal MRI artifact detection algorithm for large neuroimaging databases. However, the inference generated by DL does not commonly include a measure of uncertainty. Here, we present the first stochastic DL algorithm to generate automated, high-performing MRI artifact detection implemented on a large and imbalanced neuroimaging database. We implemented Monte Carlo dropout in a 3D AlexNet to generate probabilities and epistemic uncertainties. We then developed a method to handle class imbalance, namely data-ramping to transfer the learning by extending the dataset size and the proportion of the artifact-free data instances. We used a 34,800 scans (98% clean) dataset. At baseline, we obtained 89.3% testing accuracy (F1 = 0.230). Following the transfer learning (with data-ramping), we obtained 94.9% testing accuracy (F1 = 0.357) outperforming focal cross-entropy (92.9% testing accuracy, F1 = 0.304) incorporated for comparison at handling class imbalance. By implementing epistemic uncertainties, we improved the testing accuracy to 99.5% (F1 = 0.834), outperforming the results obtained in previous comparable studies. In addition, we estimated aleatoric uncertainties by incorporating random flips to the MRI volumes, and demonstrated that aleatoric uncertainty can be implemented as part of the pipeline. The methods we introduce enhance the efficiency of managing large databases and the exclusion of artifact images from big data analyses.

**Highlights:**

- We address the difficulty in automatically detecting artifacts in a large, imbalanced image database.
- We reproduced high accuracy in detecting artifacts in small balanced datasets.
- We implemented Monte Carlo (MC) dropout in 3D AlexNet to generate uncertainty metrics.
- We transfer the learning from a small balanced dataset to a large imbalanced dataset, outperforming focal loss.
- We outperformed artifact detection in previous studies using comparable large imbalanced datasets
- The user can select an operating point to consider manual inspection vs. artifact detection error.

## 1. Introduction

Magnetic resonance imaging (MRI^1^) is increasingly being used to detect and investigate brain morphological changes associated with neurological disorders such as Schizophrenia, Alzheimer’s disease and Parkinson’s disease (Fernandez et al., 2018; Knopman et al., 2018; Yau et al., 2018; Jubault et al., 2011). Unfortunately, MRI utility in morphometric studies is routinely compromised by image artifacts, such as those presented in Fig. 1. These artifacts can be caused by inhomogeneity of the main magnetic field, non-uniform radio frequency transmit and receive causing field bias, radio frequency spikes, subject movement, MR technician mistakes, etc. Failure to exclude volumes with such artifacts frequently causes automated morphometric analysis routines to misclassify tissue type and yield incorrect results. Such errors can be propagated into subsequent analyses involving gray matter intensity, shape, or cortical surface analysis, leading to spurious results (Ducharme et al., 2016; Reuter et al., 2015). In addition, in a multi-site study, artifacts that are specific to a particular device or site may introduce bias in the dataset. It is therefore critical to perform quality control and to screen MRI data for artifacts before using them in morphometric analyses. Moreover, timely artifact detection can aid in prompt MRI re-acquisition in longitudinal studies, especially if the number of available subjects is limited.

**Fig. 1.**
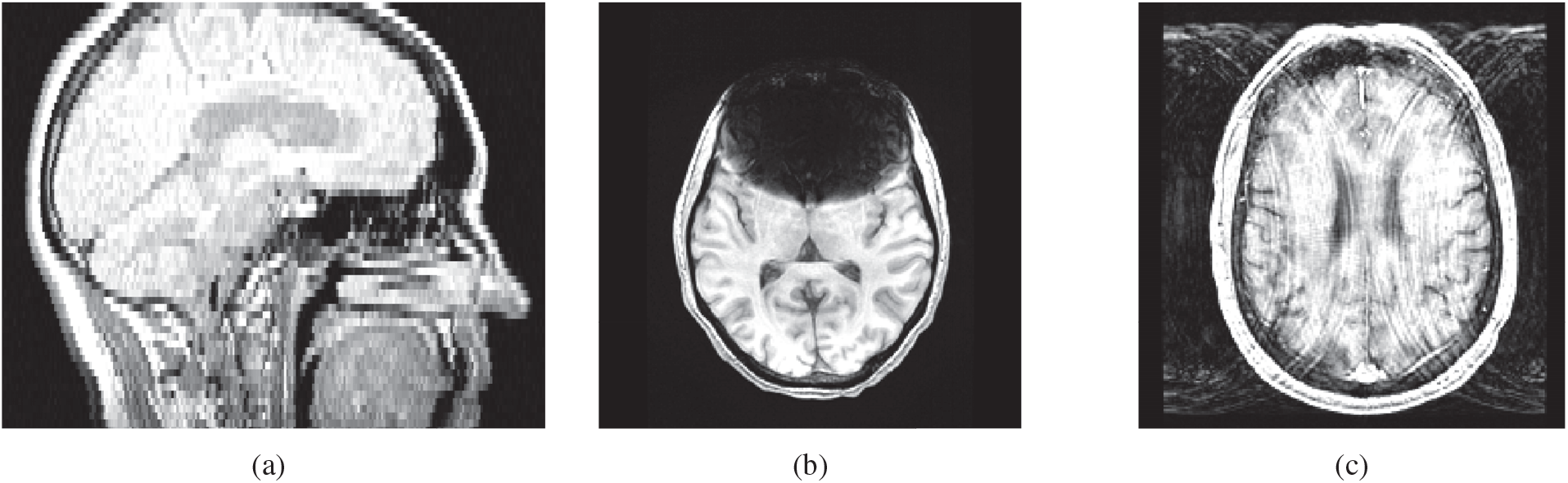
Artifact examples. The three most recurrent artifacts in our database are illustrated above: (a) coverage artifact caused by misplaced field of view, often causing wrap around, (b) intensity artifact with susceptibility caused by metallic implants, and (c) motion artifact with Gibb’s ringing caused by head movement.

Currently, the gold standard of MRI quality assessment is visual inspection by qualified personnel. However, visual inspection is a repetitive, time-consuming task, based on subjective evaluations. Indeed, Gardner et al. (1995) showed that human observers demonstrate poor sensitivity when evaluating intentionally degraded MR images, as opposed to an automated approach that can detect even minimal noise. In addition, visual inspection of MR images with subtle artifacts yields variable results. This can be demonstrated when several investigators rate MR images, as different raters can assess the same image differently. These difficulties magnify when the dataset is large and imbalanced.

In practice, datasets are typically highly imbalanced: the majority of images are of good quality. Although the primary aim of quality assessment is to identify volumes containing artifacts, ideally, the process should also retain a large proportion of the usable volumes. This brings about a repetitive and time-consuming undertaking, adding to the difficulties involved with visual inspection. These factors contribute to the challenges involved in maintaining and updating the image quality information of large and imbalanced MRI datasets in a timely manner. Automating the image quality rating could improve the reproducibility, reliability, and efficiency of image analysis, particularly when maintaining large and imbalanced neuroimaging databases (Alfaro-Almagro et al., 2018; Marcus et al., 2013; Di Martino et al., 2014).

Over the past two decades, investigators have developed automated methods to detect MRI artifacts, achieving high accuracies on small balanced datasets but only limited performance on large and imbalanced datasets. Convolutional neural network (CNN) based deep learning (DL) algorithms using patches were developed to detect motion on small testing sets of less than 20 MRI volumes (Küstner et al., 2018; Fantini et al., 2018). In another study, Oksuz (2020) used a CNN based DL algorithm to detect synthetically generated artifacts. These studies implemented state-of-the-art DL techniques and reported greater than 90% accuracies obtained with small balanced datasets; however, it remains unclear if they can generalize to large neuroimaging databases, with highly imbalanced numbers of items in different classes. Extending into a larger (> 700 MRI volumes) and highly imbalanced datasets, investigators have developed quality indices (Mortamet et al., 2009), machine learning algorithms (Pizarro et al., 2016; Alfaro-Almagro et al., 2018), binary classifiers (Esteban et al., 2017), and an ensemble DL model (Sujit et al., 2019) with limited – smaller than 90% – accuracies. To the best of our knowledge, a high-performing algorithm for artifact detection that can generalize to large imbalanced neuroimaging datasets is yet to be developed.

DL is the ideal MRI artifact detection algorithm for large neuroimaging databases, as DL works well when trained on large datasets; indeed encouraging results have been reported (Fantini et al., 2018; Küstner et al., 2018; Oksuz, 2020; Sujit et al., 2019). However, the inference generated by conventional DL algorithms does not offer a measure of prediction uncertainty, a useful metric in clinical studies; further, new data, not necessarily similar to the training set can generate misleading predictions when passed through the softmax function without an uncertainty estimation. Recently, Gal and Ghahramani (2016) developed Monte Carlo (MC) dropout, a stochastic DL algorithm that can generate inferences with probabilities and uncertainties. Several neuroimaging studies have implemented uncertainties based on MC dropout to improve lesion detection and segmentation (Nair et al., 2020; Herzog et al., 2020; Zhao et al., 2018). Uncertainties have been modeled in DL for neuroimage enhancement using a heteroscedastic model and approximate Bayesian inference (Tanno et al., 2021). Yet, to the best of our knowledge, uncertainty metrics have not been reported in studies that aim to detect MRI artifacts for large neuroimaging databases. Here we implemented a 3D AlexNet (Rani et al., 2022; Chen et al., 2021; Menon et al., 2020), to first apply a deep neural network algorithm and reproduce the high performance previously obtained with a small balanced dataset. We then develop a method to utilize this algorithm for analyzing large, imbalanced neuroimaging databases while maintaining the detection performance. We present the first stochastic DL algorithm to generate automated high-performing MRI artifact detection that can be implemented on a large and imbalanced neuroimaging database. To this end, we applied uncertainty metrics to screen volumes whose prediction confidence were relatively lower than the remaining dataset. The paper is organized as follows. In the methods, we first introduce the artifact categories, and the three stages of algorithm development through which we extended the size of the dataset and the fraction of the artifact-free data instances. Next, we describe the architecture, tuning the hyperparameters, and training methods used to obtain high performance with small datasets and large datasets. We then describe the metrics we used for evaluation after implementing MC dropout, including two artifact detection metrics developed by Alfaro-Almagro et al. (2018). We summarize our results with confusion matrices, receiver operating characteristic (ROC) plots, and tables to describe the performance of the algorithm. We conclude with a discussion.

## 2. Methods

The database we analyzed was constructed through a software pipeline to process MRI datasets derived from different neuroimaging sites contracted by companies running clinical trials or studies. The parties involved in acquiring the neuroimaging data was described in (Pizarro et al., 2019). For the present study, we curated a dataset composed of 34,800 MRI volumes, acquired from 5513 subjects in 1029 neuroimaging sites. Although there are subjects with multiple visits or longitudinal records in the full database, there are no subjects with multiple longitudinal records (or visit) in the curated dataset. When curating the dataset, we were careful to only select one visit per subject to avoid introducing potential bias in the results. A pipeline was developed for efficient processing of the data. The pipeline consisted of multiple sequential phases. MRI artifact identification is a critical task performed at the initial phase of the pipeline. Successful output of a previous phase was a prerequisite prior to processing the next phase.

In the beginning of the pipeline, MRI volumes were visually inspected to identify any existing artifact. We used the results of the visual inspection process as the ground truth for this project. We developed a DL algorithm with a 3D AlexNet to automatically identify artifacts in brain MRI volumes based on the ground truth. To that end, the CNN architecture was developed by tuning the hyperparameters, incorporating MC dropout, and identifying an unbiased set of performance metrics. The algorithm was developed over three successive stages allowing us to (I) test the proof-of-concept, extend to (II) an intermediate balanced dataset, and (III) a realistic large imbalanced dataset.

### 2.1. Curating a well labeled dataset

Building on our success in developing DL algorithms for MRI (Pizarro et al., 2019), one of our main challenges in this project was curating a well-defined dataset. Due to the nature of this project, identifying a dataset was not straightforward given the common challenges faced in the field. We encountered DL challenges, such as, data heterogeneity (Yu et al., 2019), shortage of diverse samples, data quality, and data dependency (Munappy et al., 2019). In addition, we encountered Big Data challenges such as discriminative tasks, high-dimensional data, and extracting complex patterns from massive volumes of data (Najafabadi et al., 2015). We overcame these challenges and compiled a curated dataset to train and test the proposed DL algorithm.

#### 2.1.1. Identifying MRI with artifacts

In our experience, properly labeling MRI volumes with artifacts is challenging and time consuming, thus limiting the size of the curated dataset. To identify and classify the three most recurrent artifacts in our database, we used the LORIS guidelines (Das et al., 2012). Table 2 summarizes the criteria, the corresponding manifestation, and possible causes of the artifacts. MRI volumes in our study belonged to one of seven structural MRI contrasts as detailed in (Pizarro et al., 2019), with the following exclusions:

**Table 1.** Abbreviations used throughout manuscript.

| Abbreviation | Meaning |
| --- | --- |
| AUC | area under the curve |
| CNN | convolutional neural network |
| CP | clean percentage |
| DFFMR | datasets flagged for manual review,<br>i.e., visual inspection |
| DL | deep learning |
| LR | learning rate |
| MRI | magnetic resonance imaging |
| MC | Monte Carlo |
| MI | mutual information |
| ROC | receiver operating characteristic |
| UDM | unusable datasets missed |

**Table 2.** The three most recurrent artifacts in our database: class, type, and cause.

| artifact class | artifact type | cause |
| --- | --- | --- |
| intensity | hyper-intensity | bias (receive/transmit) fields |
|  | signal drop-off | RF coil, misplaced saturation band |
|  | susceptibility | implants and fashion accessories |
| motion | ghosting | head movement |
|  | banding aka ‘corduroy’ artifact | head movement |
|  | Gibbs ringing | head movement |
| coverage | wrap around or aliasing | misplaced field of view |
|  | part of brain is cut | misplaced field of view |

- MRI volumes acquired on older MRI scanner models that are no longer being used.
- MRI volumes acquired with parameters that were significantly different from the standard ones (e.g., resolution, pulse sequence type).

We defined an artifact as a misrepresentation of tissue structures produced by the acquisition sequence, which prevents meaningful image processing or analysis. The artifact dataset in all three stages consisted of MRI volumes containing prominent artifacts. The MRI volumes with artifacts used in this study failed the quality control checks performed within the context of a clinical trial. As part of quality control, the users of the database were trained to identify image artifacts and, if necessary, fail the data obtained in the subject’s MRI session. In this study, we aimed to develop automated tests to detect artifacts that are usually detected by careful visual inspections and cannot be detected by examining the image metadata. To this end, we collaborated with database managers who defined a comprehensive list of quality control tests. Among other tests, these quality control tests included checking the acquisition parameters, the visibility of the contrast agent (typically Gadolinium), and visual inspection of the images. These quality control tests were previously applied to the images in the large database we have used. The dataset available for training had unstructured text descriptions of the reason for failure. This text included imaging but also non-imaging-related issues, and it covered multiple scans from the same session. To develop our automated tests, we narrowed our search for the MRI volumes with artifacts by using all the images from sessions in which at least one image failed these quality control checks. We visually inspected the MRI volumes that had previously failed quality control checks to curate the MRI volumes with visually visible artifacts.

We visually inspected each MRI volume that had ‘failed’ quality control, to generate a list of MRI volumes that contained at least one artifact class according to Table 2. Visually inspected scans contained either no artifact, one type of artifact, or more than one type of artifact. Albeit a subjective process, we quantified the level of visible artifact present in the image with explanatory terms: no visible artifact, mild artifact, and prominent artifact. We included in the artifact dataset images that were labeled to have at least one prominent artifact. The artifact dataset consisted of 535 MRI volumes in stage I and 696 MRI volumes in stages II and III (in stages II and III we incorporated additional MRI volumes that had been acquired after stage I completion). The goal of the algorithm we present is to accurately identify MRI volumes with artifacts. Database managers are rarely interested in distinguishing among artifacts of different types; rather, they are primarily concerned with screening out MRI volumes with artifacts, in order to avoid propagating problems in the processing and analysis pipeline. We initially trained on more than two classes but found that using only two classes was beneficial for the following reasons. First, using only two classes generated higher accuracies in the testing phase. Second, we addressed the difficulties associated with detecting artifacts in large, imbalanced datasets by applying data ramping combined with transfer learning. Using multiple classes would make it more difficult to correctly split the imbalanced datasets.

#### 2.1.2. Incorporating clean MRI volumes

We paired the artifact MRI dataset with a clean – artifact-free – MRI dataset. To develop the algorithm along each stage, we increased the size of the dataset and the clean percentage (CP), i.e., the fraction of the artifact-free MRI volumes. For proof-of-concept in stage I, we visually inspected a set of images and identified a list of 101 clean MRI volumes, creating a dataset with CP=15.9%. The clean MRI volumes from stage I met two criteria: (i) they had passed the initial quality control tests including the visual inspection, and they had been successfully processed with the pipeline developed in-house, i.e., had not been labeled ‘failed’ and (ii) Based on an additional visual inspection, the MRI volumes were visually free from any artifact. Stage I results showed that it was possible to implement DL to distinguish MRI volumes with artifacts from clean MRI volumes. However, the stage I dataset was not realistic, as the clean images did not represent the entire database due to two issues:

A. The clean images accounted for only 15.9% of the curated dataset, CP=15.9%.
B. The artifact dataset were obtained in a set of studies, **s**_*a*_, while the clean dataset was taken from a different study **s**_*c*_, such that **s**_*a*_ ∩ **s**_*c*_ = ∅.

To address issue A, we incorporated hundreds of additional images from our database, to create a more representative sample of clean images, that have been visually inspected by MRI experts. In stages II and III, we identified MRI volumes to be clean if they met criterion (i), without the additional inspection, namely criterion (ii). With this criterion, we quickly and efficiently increased the clean percentage in the dataset: CP=50% in stage II, and realistic CP=98% in stage III. To address issue B, in stages II and III we matched the distribution of clean datasets obtained in different studies to the corresponding distribution of the artifact datasets, by keeping the CP approximately constant across studies. Some MRI volumes with artifacts were acquired in studies that did not have clean MRI volumes available for analysis. We incorporated these constraints to develop a process entitled Clean Selector, illustrated in Supplementary Section Appendix A, and used in stages II and III. After successfully incorporating clean MRI volumes for each stage, we tested the generalizability of the algorithm on unseen data, by applying cross-validation and splitting the dataset as shown in Table 3.

**Table 3.** Data for cross validation for stages I, II, and III. The used datasets contained MRI volumes with and without artifacts. Abbreviations: MRI - magnetic resonance imaging and CP - clean percent.

| cross validation subset | split (%) | MRI volumes | clean | artifact | CP |
| --- | --- | --- | --- | --- | --- |
| training | 75 | 477 | 76 | 401 | 15.9% |
| validation | 5 | 32 | 5 | 27 | 15.6% |
| testing | 20 | 127 | 20 | 107 | 15.7% |
| **stage I proof-of-concept** | **100** | **636** | **101** | **535** | **15.9%** |
| training | 60 | 834 | 417 | 417 | 50.0% |
| validation | 20 | 279 | 140 | 139 | 50.2% |
| testing | 20 | 279 | 139 | 140 | 49.8% |
| **stage II balanced** | **100** | **1392** | **696** | **696** | **50.0%** |
| training | 60 | 20880 | 20463 | 417 | 98.0% |
| validation | 20 | 6960 | 6821 | 139 | 98.0% |
| testing | 20 | 6960 | 6820 | 140 | 98.0% |
| **stage III realistic** | **100** | **34800** | **34104** | **696** | **98.0%** |

### 2.2. Deep learning with convolutional neural networks

We used our labeled dataset to develop a DL algorithm with a 3D AlexNet to infer whether a brain MRI intensity image had an artifact. That is, we used image data instead of using header files, global metrics (SNR, CNR) or other metrics derived from image histogram, as in (Pizarro et al., 2019). The input to the DL algorithm mimics what human-rater sees when viewing an MRI volume. We developed the algorithm in Python with a Tensorflow backend and compiled on Keras (Chollet, 2017); the implementation is available on GitHub (a link will be added after the manuscript is accepted for publication). The DL algorithm was compiled to run on a Nvidia Quadro K2200. To minimize complexity and overfitting, the architecture hyperparameters were selected empirically, except for the learning rate (LR), which was systematically tuned to incorporate cyclical LR (Smith, 2017).

#### 2.2.1. Convolutional neural network architecture

We designed several distinct networks by modifying the architecture hyperparameters, including: the number of layers, the kernel dimension, the stride dimension, the number of filters (or nodes) by layers, normalization (batch or instance), and convolutional padding convention (same or valid). We applied dropout in layers 17 and 19 (in Fig. 2) during training for all three developmental stages. In stages II and III we incorporated MC dropout by coding the corresponding layers to also perform dropout during inference of the validation and testing subsets. We empirically selected the 3D AlexNet architecture in Fig. 2 by striking a balance between complexity and overfitting, while optimizing performance. During training, we performed image intensity normalization prior to using the images as input to the 3D CNN. We performed image intensity normalization by subtracting the mean image intensity and dividing by the standard deviation of the image intensity, as we did in (Pizarro et al., 2019). We then paired the input, *X* – image intensity normalized MRI volumes with 256 × 256 × 64 dimensions (voxels) – with the output, *Y* – the labeled ground truth, *l* ∈ {0, 1}, with 2 × 1 dimensions, i.e., clean or artifact. The layers type and description were fully described in (Pizarro et al., 2019), with one difference: here we incorporated convolution in 3-dimensions instead of 2-dimensions.

**Fig. 2.**
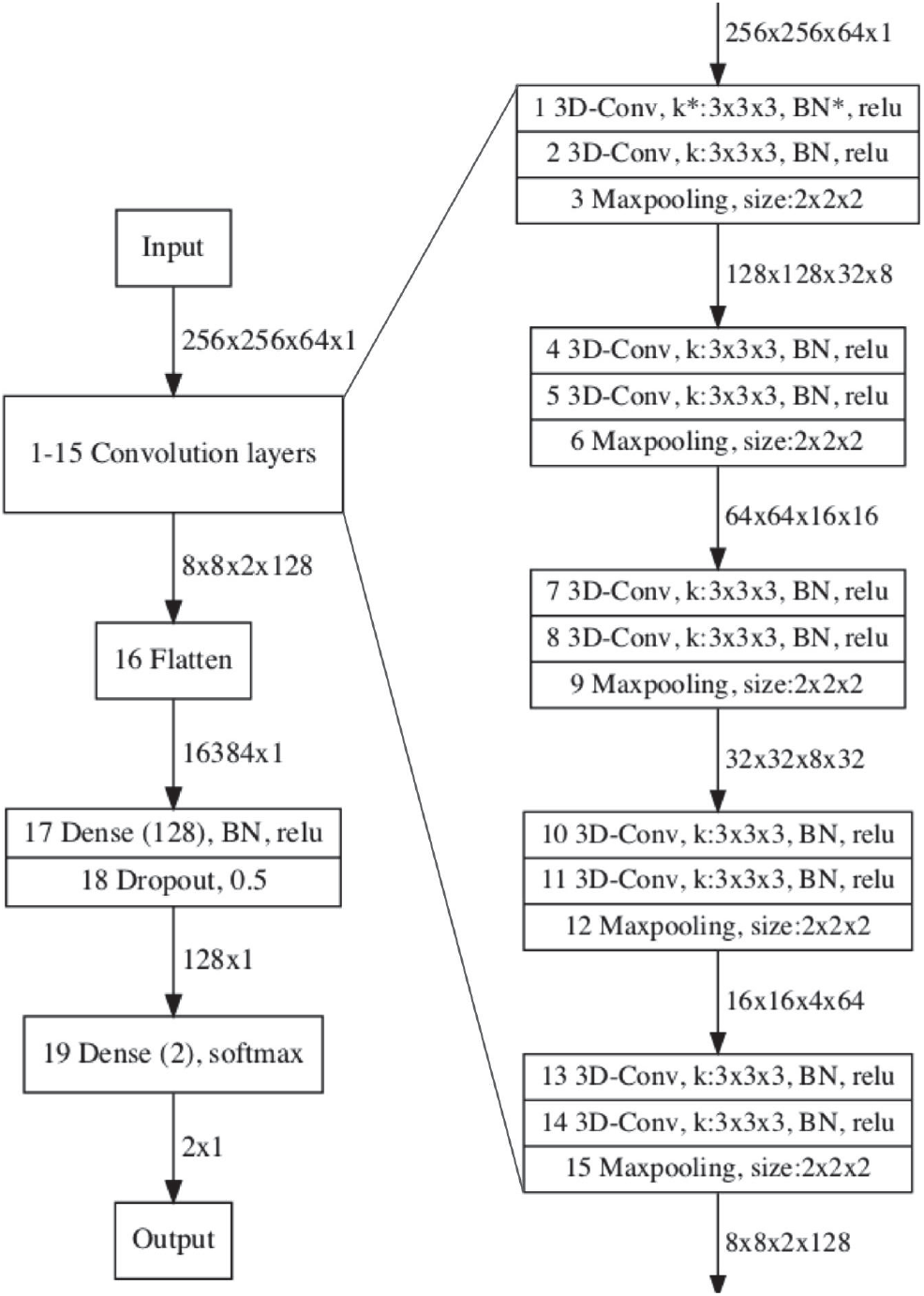
Three-dimensional AlexNet (Rani et al., 2022; Chen et al., 2021; Menon et al., 2020) architecture used for deep learning. The input was a single MRI volume, and the output generated an inference of two categories: clean or artifact. Abbreviations: MRI - magnetic resonance imaging, 3D-Conv - 3D convolution, k - kernel dimension, BN - batch normalization, and relu - rectified linear unit.

For this project, the image processing steps involved commonly used techniques to prepare the data for DL, similar to those we applied in (Pizarro et al., 2019). For each MRI volume, we performed intensity normalization and resized the image data to fit into the dimensions specified in the architecture. Our goal was to develop an automated algorithm for detecting artifacts that can be applied directly to the incoming MRI data before sending them through the processing pipeline. An additional processing step at the beginning of the pipeline, would result in another potential point of failure and increased chances for errors. Simplifying the automatic detection of artifacts by using native images directly has the potential to further improve the pipeline’s reliability and generalizability.

#### 2.2.2. Tuning the learning rate

During training, we incorporated cyclical LR (Smith, 2017) by varying between reasonable values, found by employing two LR range tests. The LR range test – estimating performance with changing LRs – is highly valuable whenever using a new architecture or dataset. To that end, we conducted two LR experiments, to identify the LR that generated the fastest decrease in the loss. Then during training, we defined a range *R*_*T*_, around this optimal point to incorporate cyclical LR for efficient optimization. We trained the DL algorithm for *E* epochs. For efficiency, we folded the data into *f* parts and sequentially rotated over each part, composed of *s* _*f*_ scans. This resulted in using each of the *f* parts to train for *E*/*f* epochs. In experiment 1, MRI volumes with artifacts were introduced first, while in experiment 2, the two classes were interspersed. We exponentially increased the LR for each epoch spanning the range *R*_*exp*_, for each experiment. Each epoch took δ steps and used *n* MRI volumes per step. The parameters are summarized in Table 4. We tracked the loss (categorical cross-entropy) over *E* epochs and plotted versus the LR on a log-scale in Fig. 3. Next, we computed a differential loss (*dloss*) and used a moving average with a window of *w* units to plot *dloss* versus LR in Fig. 3. The LR that minimized *dloss* is denoted with a red star.

**Table 4.**
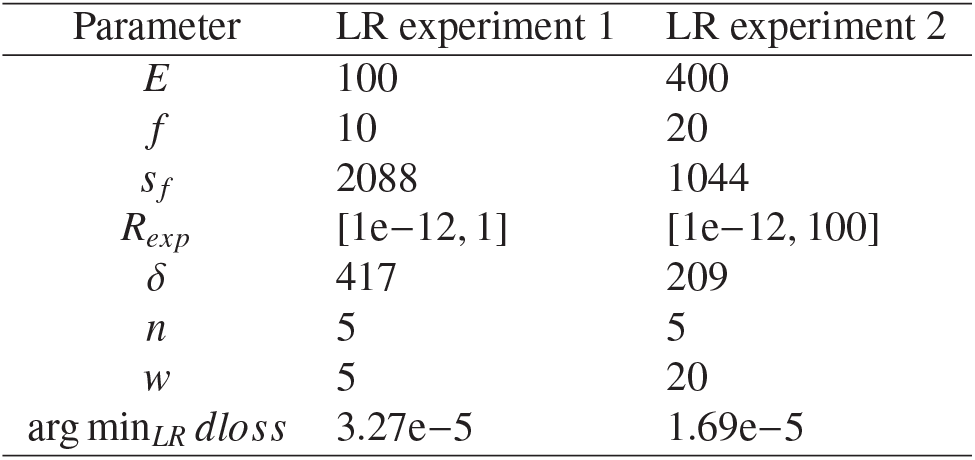
Parameters used for each LR experiment. Abbreviations: LR - learning rate, *E* - number of epochs, *f* - number of folds, *s* _*f*_ - scans per fold and epoch, *R*_*exp*_ - LR range for experiments, δ - steps per epochs, *n* - MRI volumes per step, *w* - moving average window, and arg min_*LR*_ *dloss* - LR that minimized differential loss.

**Fig. 3.**
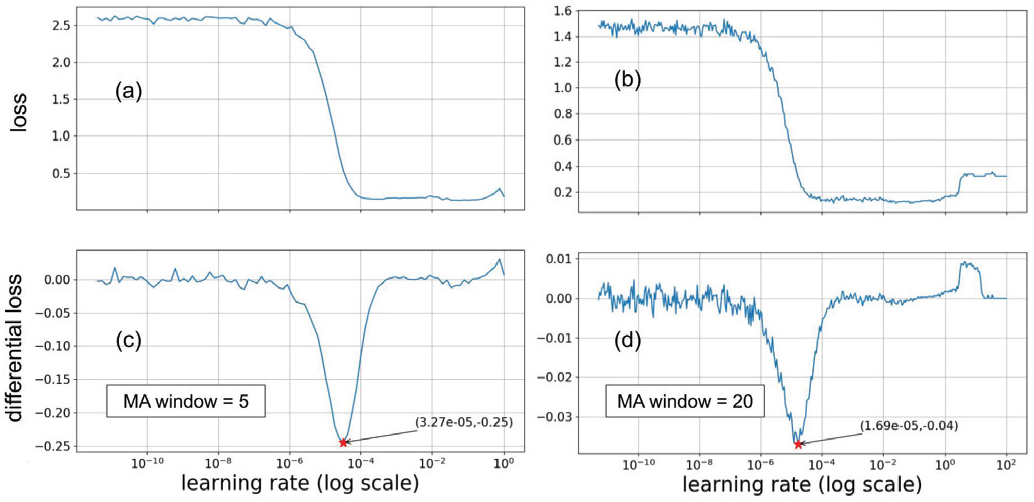
Tuning the LR. We ran two experiments to tune to LR by tracking the categorical cross-entropy (loss) as a function of LR (log-scale) (upper row, (a) and (b)). We computed the differential loss, then smoothed with an MA window, and plotted as a function of LR (log-scale) (bottom row, (c) and (d)). The LR that minimized the differential loss is denoted by a red star, where the loss decreased at the fastest rate. Abbreviations: LR - learning rate and MA - moving average.

**Fig. 4.**
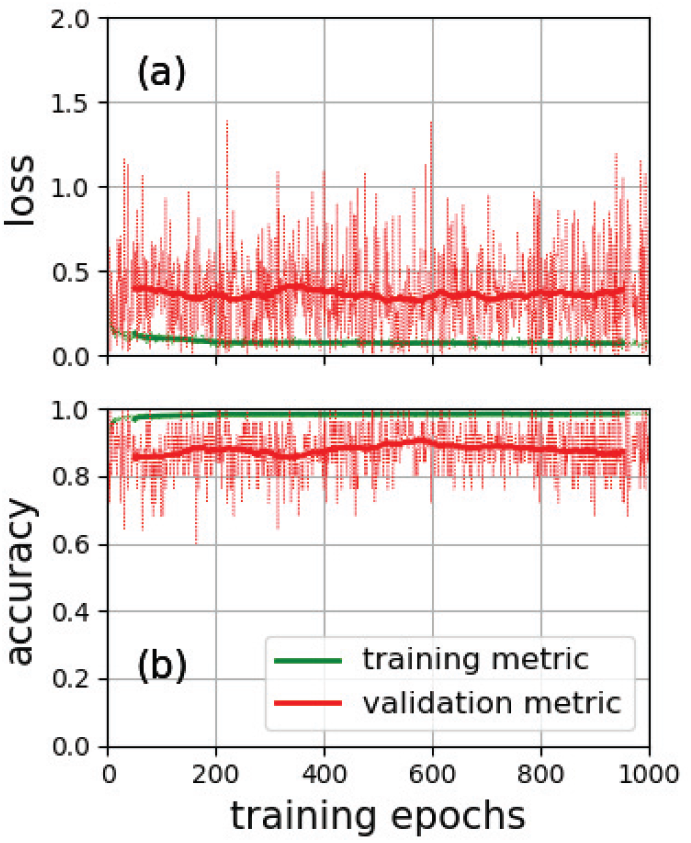
Stage I performance metrics. The metrics obtained in 1000 training epochs are plotted for the training (in green) and validation (in red) subsets. The loss (categorical cross-entropy) is plotted in (a) and the accuracy in (b). For each metric the value is plotted with a dashed line at the current epoch along with a moving average with a solid line.

### 2.3. Development, implementation, and training

After developing the architecture along with tuning the LR, we went on to developing the three stages with increasing dataset size and percentage of clean MRI volumes, as detailed in Table 3. The algorithm was trained to minimize the loss, defined to be the categorical cross-entropy (Pizarro et al., 2019). While training, we tracked the loss and accuracy for both the training and validation subsets, where accuracy is the proportion of correctly identified to total MRI volumes in the subset. In stage I, we used a small dataset with CP=15.9%, to test the proof-of-concept. In stage II, the dataset was balanced to CP=50%, and used to efficiently develop a high performing algorithm by implementing probability and uncertainties. In stage III, the dataset was extended to a realistic CP=98% and used to develop a training process to transfer the learning from stage II. After the training completed in each stage, the model parameters were used to predict the data from corresponding validation and testing subsets. In addition, the model parameters saved after the completion of stage II were used for initializing four of the five stage III experiments, as specified in Section 2.3.3. To demonstrate the complexity of developing a high performing algorithm for large imbalanced datasets, we included (in Figures 6 and 8) the results of two unsuccessful experiments.

**Fig. 5.**
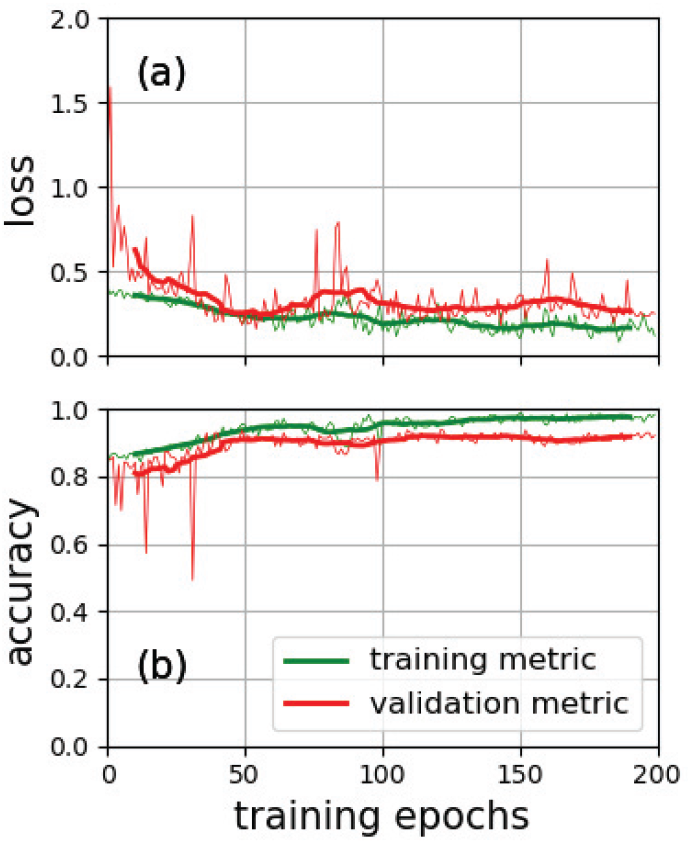
Stage I performance metrics after applying ComBat Fortin et al. (2018) to the proof-of-concept dataset. The metrics obtained in 200 training epochs are plotted for the training (in green) and validation (in red) subsets. The loss (categorical cross-entropy) is plotted in (a) and the accuracy in (b). For each metric the value is plotted with a dashed line at the current epoch along with a moving average with a solid line.

**Fig. 6.**
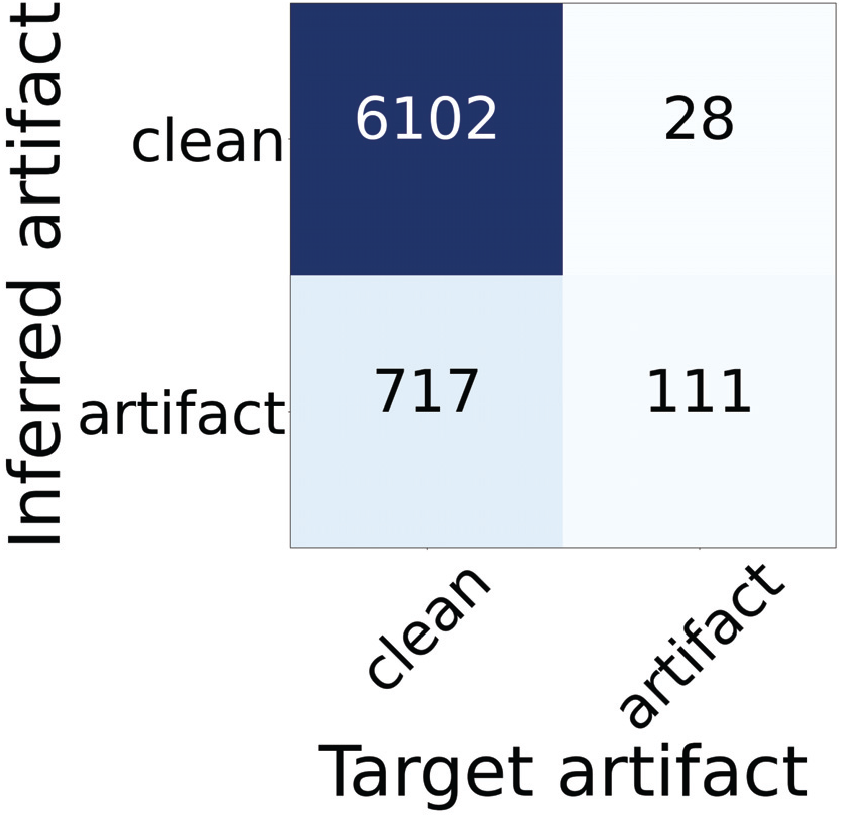
Confusion matrix for an initial attempt to classify a realistic dataset with CP=98%. The target was taken from the ground truth label obtained by visual inspection. The inferred artifact was determined using the proposed algorithm. The predictions are summarized as F1 = 0.230, ACC = 0.893, SPC = 0.895, TN = 6102, FP = 717, SEN = 0.799, TP = 111, and FN = 28. Abbreviations: ACC - accuracy, SPC - specificity, TN - true negative, FP - false positive, SEN - sensitivity, FN - false negative, TP - true positive. Note that negative stands for clean MRI volume (usable), and positive stands for artifact MRI volume (unusable).

**Fig. 7.**
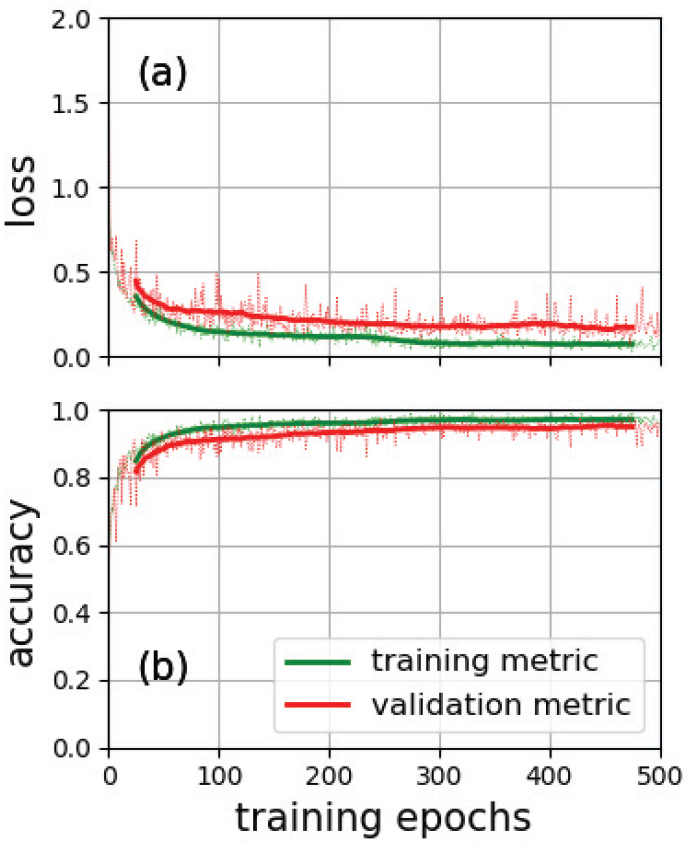
Stage II performance metrics. The metrics obtained in 500 training epochs are plotted for the training (in green) and validation (in red) subsets. The loss (categorical cross-entropy) is plotted in (a) and the accuracy in (b). For each metric the value is plotted with a dashed line at the current epoch along with a moving average plotted with a solid line.

**Fig. 8.**
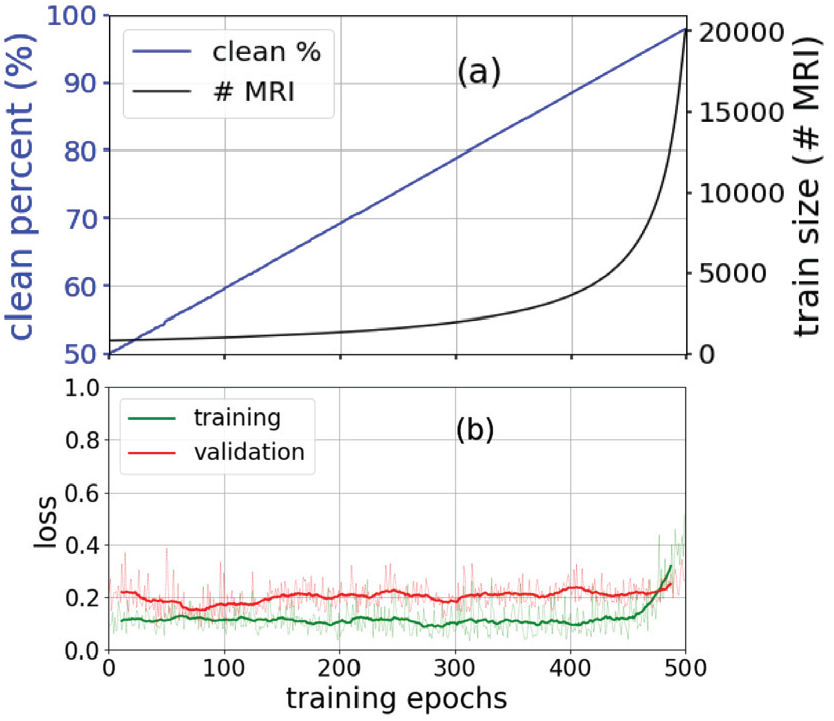
Clean percentage ramping experiment. (a) The clean percentage ramping experiment was designed to linearly ramp the clean percentage (blue) from 50% to 98% over 500 epochs, which resulted in asymptotically increasing the train set size (black). (b) The loss (categorical cross-entropy) generated by the training (green) and validation (red) subsets along with a moving average.

**Fig. 9.**
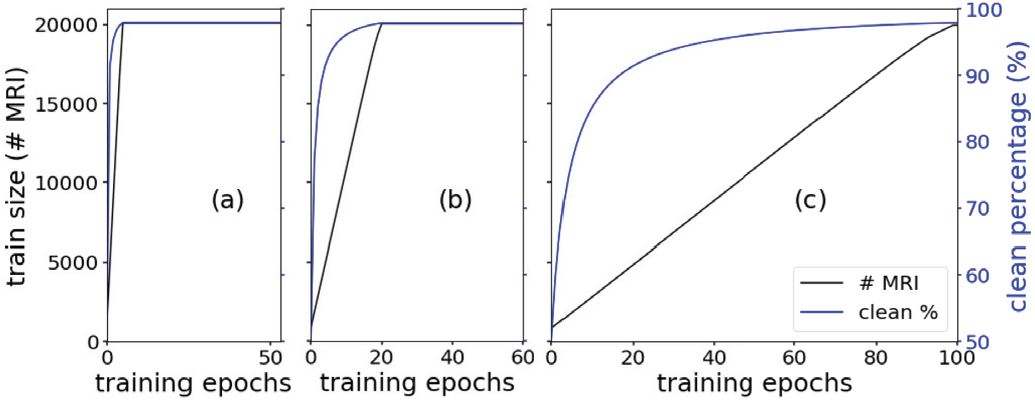
Design for data-ramping experiments. We linearly ramped the data from 834 MRI scans (50% clean) up to 20,880 MRI scans (98% clean), over three different ramping epochs, *E*_*R*_ = {5, 20, 100}, respectively plotted in (a), (b), and (c). The dataset size is plotted in black (scale and units on the left vertical axis) alongside the corresponding clean percentage in blue (scale and units on the right). The three values of *E*_*R*_ resulted in training the data-ramping experiments for *E* = {53, 60, 100} epochs.

#### 2.3.1. Stage I: Testing the proof-of-concept with a small dataset

We implemented a previously developed training process (Pizarro et al., 2019) with the small proof-of-concept dataset (Table 3). We initialized with random parameters. Each epoch used 500 estimation steps and a batch size of 5. The training process used 2500 volumes per epoch for 1000 epochs. The loss (categorical cross-entropy) and accuracy are plotted in Fig. 4 for the training and validation subsets. After the training completed, the model parameters were used to predict the data from the testing subsets; the results are summarized in Fig. 15.

To address distribution differences between clean and artifact-labeled datasets used in stage I, we investigated domain adaptation methods (de Mathelin et al., 2021). Fortin et al. (2018) showed that variance of non-biological origin is introduced by differences in MRI scanners and acquisition protocols. Therefore, having the training and validation sets come from different studies (different clinical trials) may have boosted the accuracy we obtained in Stage I before applying domain adaptation. We implemented ComBat which was developed to harmonize the data across MRI neuroimaging sites (Fortin et al., 2018). We hypothesized that applying ComBat for domain adaptation to the dataset used in stage I would result in lower testing accuracy. We applied neuroCombat (https://github.com/Jfortin1/neuroCombat) to our dataset, by selecting the study (i.e., the clinical trial) as the scaling factor. That is, we harmonized the proof-of-concept dataset in stage I across the clean and artifact-labeled datasets. To meet the computational resources required for neuroComBat, neuroCombat was implemented in batches. We divided the entire stage I (proof-of-concept) dataset of 636 MRIs (101 clean MRI and 535 artifact MRI data sets) into four sub-sets of 159 each and applied neuroCombat harmonization. Then, we split the harmonized dataset into its corresponding cross-validation subsets and retrained the 3D AlexNet algorithm. The loss (categorical cross-entropy) and accuracy are plotted in Fig. 5 for the training and validation subsets. After the training was completed, the model parameters were used to generate inferences on the data from the testing subsets; the results are summarized in Fig. 16.

#### 2.3.2. Stage II: Making the dataset balanced and developing uncertainties

After obtaining encouraging results on an unseen testing subset in stage I, we tried using a realistic dataset, with CP=98%. We trained the algorithm with a similar process used in stage I (details and performance metrics not presented) then generated inferences on an unseen testing subset, presented in Fig 6. We summarized the performance with accuracy = 0.893 (F1-score = 0.230, specificity = 0.895 and sensitivity = 0.799). The unimpressive results made it obvious that we needed to continue developing the algorithm. Further, training the algorithm with the realistic dataset was time consuming, motivating the creation of stage II for training based on the balanced dataset. We used the balanced dataset (CP=50%) and adapted the stage I training process, as follows. To generate probabilities and uncertainties, we incorporated MC dropout (Gal and Ghahramani, 2016). We initialized with random parameters and trained with all 834 MRI volumes of the balanced dataset, CP=50% (Table 3), for 500 epochs, where each epoch consisted of 166 estimation steps and a batch of 5. We used the results from the LR experiments to implement a sawtooth cyclical LR (Smith, 2017) with range *R*_*T*_, and limits [*L*_*LR*_, *U*_*LR*_] fixed for the first 250 epochs at *L*_*LR*_ = 1e−6 and *U*_*LR*_ = 1e−3. During the second 250 epochs, we decreased *U*_*LR*_ exponentially by λ = 0.946 down to *L*_*LR*_. Fig. 7 presents the loss (categorical cross-entropy) and accuracy, plotted for the training and validation subsets. After training completed, the model parameters were used to predict the data from corresponding validation and testing subsets. The results are presented as ROC curves (Fig. 17) and summary statistics (Table 6).

#### 2.3.3. Stage III: Implementing transfer learning for classifying a realistic – 98% clean – dataset

Learning from our initial unsuccessful attempts of classifying a realistic dataset, we created ramping experiments to investigate methods for transferring the high-performing results obtained with a balanced dataset in stage II. Our first attempt at transferring the learning was to ramp the clean percentage of the dataset while training the algorithm, as illustrated in Fig. 8. We first used the high-performing model saved from stage II to initialize the training. Then, as the stage II dataset was balanced, we linearly ramped the clean percentage from CP=50% up to CP=98% over 500 training epochs. Prior to starting each epoch, we incorporated clean scans to increase the clean percentage by (98%-50%)/500 = 0.096%. A linear increase in clean percentage resulted in an asymptotic increase in dataset size. The clean percentage ramping experiment generated lower performance than expected. Based on the results, we hypothesized that the nonlinear increase in dataset size resulted in a similar nonlinear increase in the training loss, observed in Fig. 8. Introducing the model in the final epochs with so many new scans, albeit clean scans, resulted in the optimization unable to keep up, hence the loss increase. Regardless, we included our attempt here, to highlight how our naive approach informed and motivated the next step.

To test our hypothesis and avoid the nonlinear increase in the loss, we focused on linearly ramping the dataset size in the next set of experiments. We designed five experiments: two baseline experiments and three data-ramping experiments. For comparison under similar conditions, the time of each experiment was constrained to τ = 5 days. The first baseline experiment, entitled `S3_base_random`, was initialized with random parameters. The second, entitled `S3_base_initialized`, was initialized with the model parameters saved after stage II completion. In the two baseline experiments, we trained for *E*_*B*_ = 50 baseline epochs with the realistic training subset (Table 3) comprising *T*_*r*_ = 20880 MRI volumes (CP = 98%). The time τ, it took for the training to complete is linearly related to the amount of data used during training:

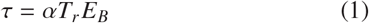

where α has units of *days*/(*MRI* · *epoch*) and is related to the speed of the graphics processing unit. We designed data-ramping experiments (Fig. 9) for linearly increasing the dataset size during training from balanced training size (*T*_*b*_ = 834 MRI scans; CP = 50%) up to realistic training size (*T*_*r*_ MRI scans; CP=98%). Each data-ramping experiment was initialized with the model parameters saved after stage II completion. We varied the ramp slope by assigning to each experiment one of three different ramping epochs, *E*_*R*_ = {5, 20, 100}, respectively entitled `S3_dr_ER_005`, `S3_dr_ER_020`, and `S3_dr_ER_100`. The ramp slope is defined by the ramping epoch *E*_*R*_, specifying the number of epochs to linearly ramp the data from *T*_*b*_ up to *T*_*r*_. The time τ, to train each data-ramping experiment, depended on the two training sizes *T*_*b*_, *T*_*r*_, the number of training epochs, *E*, and ramping epochs *E*_*R*_:

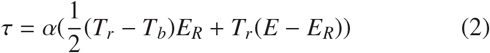

Setting Eq. 1 equal to Eq. 2, we estimated the number of training epochs *E* as a function of baseline epochs *E*_*B*_, and ramping epochs *E*_*R*_:

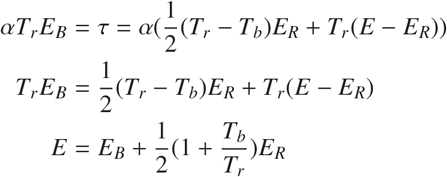

We rounded the values and trained for *E* = {53, 60, 100} epochs, respectively, ensuring each experiment completed in the same allotted time, τ. For evaluating reproducibility, we repeated the following experiments two additional times: `S3_base_initialized`, `S3_dr_ER_005`, and `S3_dr_ER_100`. For each repetition, we appended the experiment name with ‘N2’ and ‘N3’ in Section 3 to indicate that we ran the experiment a second, and third time, respectively. The additional repeats resulted in a total of 11 experiments. The experiments were designed to answer the following experimental questions:

Q1 Does the stage II model provide a good starting point to initialize training in stage III? This was addressed by comparing the results of `S3_base_random` to `S3_base_initialized`.
Q2 Does data ramping provide a path for transfer learning from stage II to stage III? This was addressed by comparing the results of `S3_base_initialized` to the three data ramping experiments: `S3_dr_ER_005`, `S3_dr_ER_020`, and `S3_dr_ER 100`.
Q3 Does the data ramp slope influence the training performance? This was addressed by comparing amongst the data ramping experiments: `S3_dr_ER_005`, `S3_dr_ER_020`, and `S3_dr_ER_100`.

As in stage II, we used MC dropout along with a sawtooth cyclical LR (Smith, 2017) with range *R*_*T*_, and limits [*L*_*LR*_, *U*_*LR*_] fixed for the first *E* − 10 epochs at *L*_*LR*_ = 1e−6 and *U*_*LR*_ = 1e−3. During the last 10 epochs, we decreased *U*_*LR*_ exponentially by λ = 0.252 down to *L*_*LR*_. In addition, we incorporated Keras’ (Chollet, 2017) ‘class weights’ to ensure the classes contributed equally to the loss (Japkowicz and Stephen, 2002). We estimated the performance of the algorithm by using the realistic validation subset (Table 3; 6960 MRI scans; CP=98%) after each epoch was completed. Fig. 10 presents the performance metrics for the two baseline experiments. Fig. 11, presents the performance metrics for the three data-ramping experiments. After training completed, the model parameters were used to predict the data from corresponding validation and testing subsets, with results presented as ROC curves (Figs. 18 and 19) and summary statistics (Table 7).

**Fig. 10.**
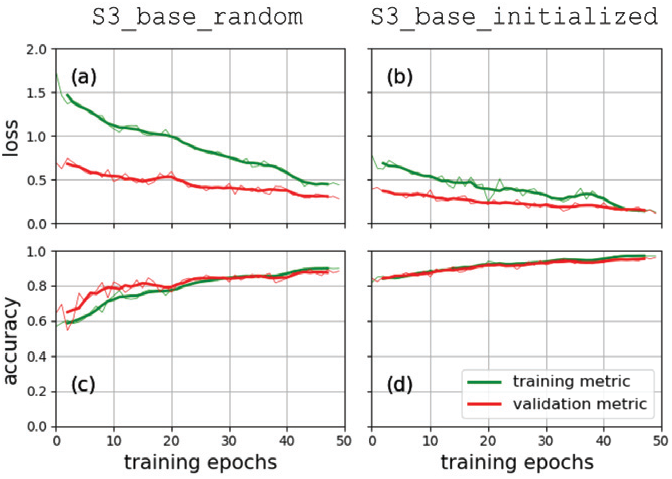
Performance metrics for the two stage III ‘baseline’ experiments. On the left ((a) and (c)), the performance is plotted for `S3_base_random` that was initialized with a random set of parameters. On the right ((b) and (d)), the performance is plotted for `S3_base_initialized` that was initialized with the model parameters saved from stage II. The loss (categorical cross-entropy) is plotted on the first row (a) and (b), and the accuracy on the second row (c) and (d). Each metric is plotted in a dashed line along with a moving average in a solid line. The training and validation performances are plotted in green and red, respectively.

**Fig. 11.**
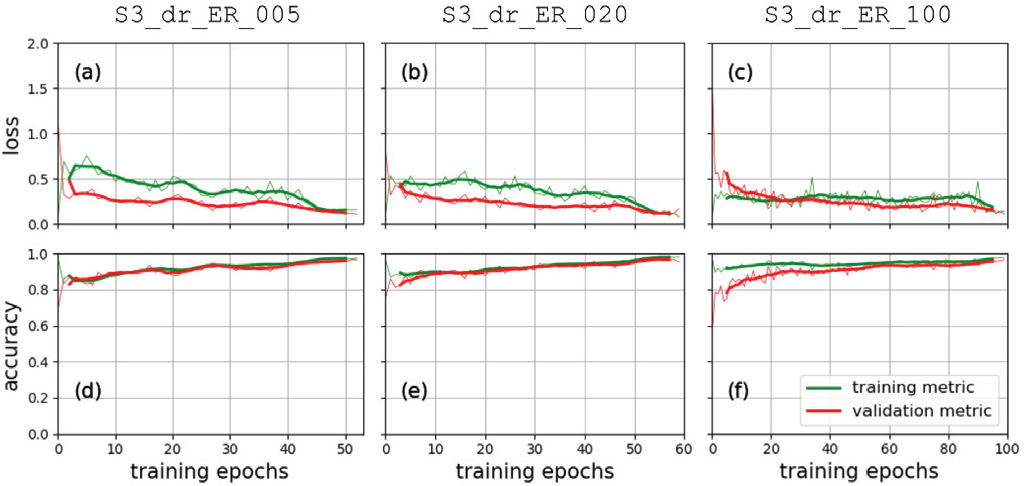
Performance metrics for the three stage III ‘data-ramping’ experiments. On the left ((a) and (d)), the performance is plotted for `S3_dr_ER_005`; in the middle ((b) and (e)), for `S3_dr_ER_020`; and on the right ((c) and (f)), for `S3_dr_ER_100`. The ramping epochs for the three experiments were *E*_*R*_ = {5, 20, 100}, respectively. The loss (categorical cross-entropy) is plotted on the first row (a)-(c), and the accuracy on the second row (d)-(f). Each metric is plotted in a dashed line along with a moving average in a solid line. The metrics for the training and validation performances are plotted in green and red, respectively.

**Fig. 12.**
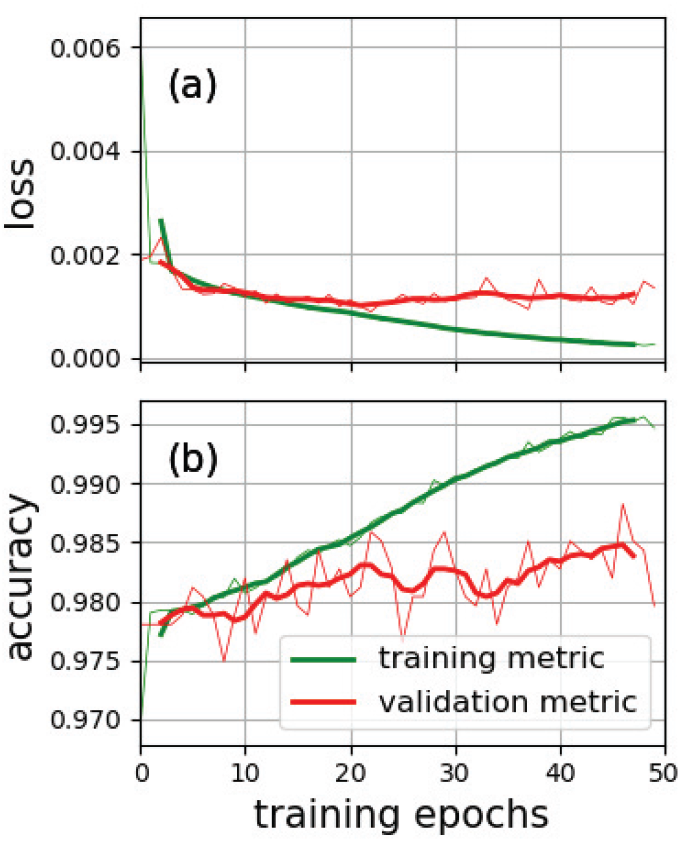
Performance metrics for the S3 focal loss experiment. The metrics obtained in 50 training epochs are plotted for the training (in green) and validation (in red) subsets. The loss (focal cross-entropy) is plotted in (a) and the accuracy in (b). For each metric the value is plotted with a dashed line at the current epoch along with a moving average plotted with a solid line. Note: The range of the y-axis for loss and accuracy was selected to provide informative performance plots.

**Fig. 13.**
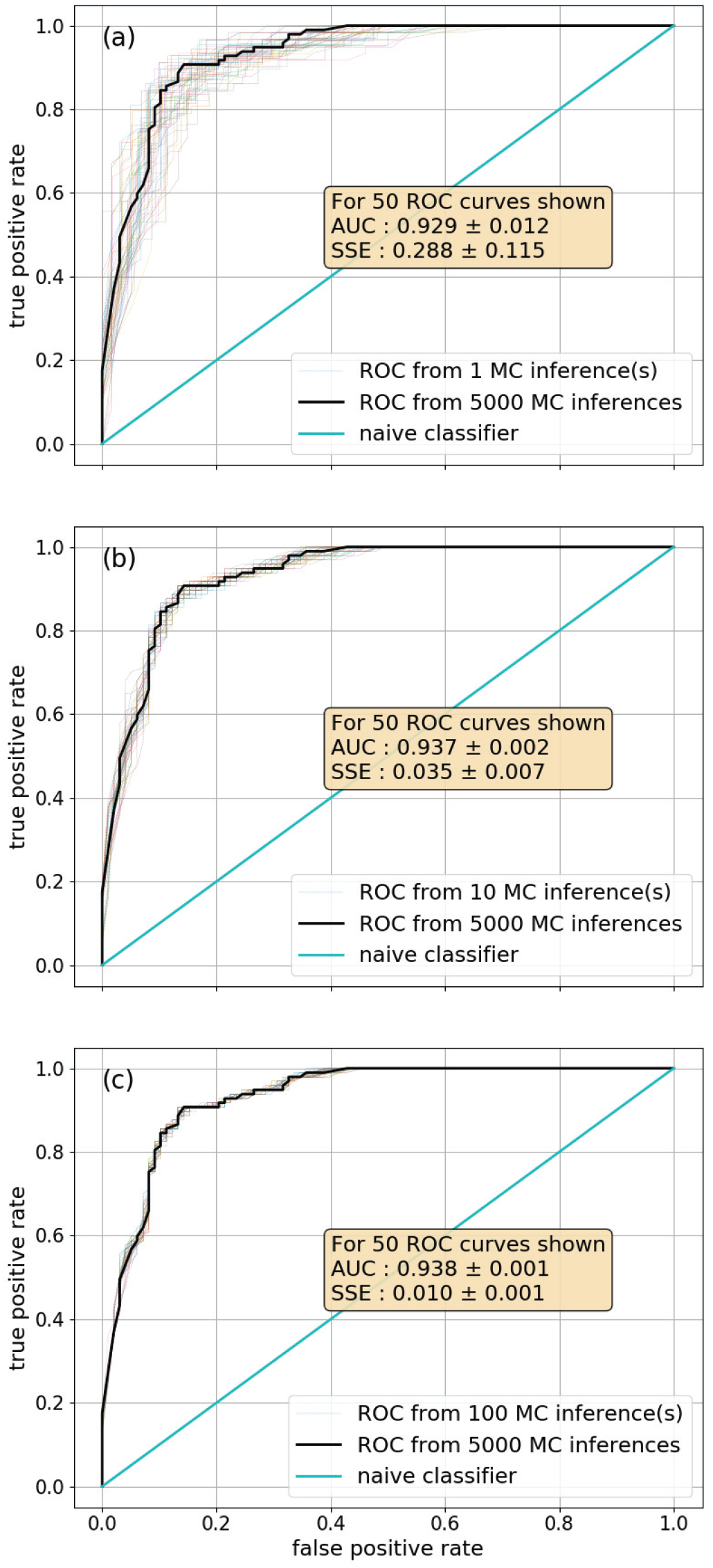
Estimating the number of MC inferences needed. Fifty ROC curves were generated from MC dropout inferences by averaging candidate values (a) *N*_*C*_ = 1, (b) *N*_*C*_ = 10, and (c) *N*_*C*_ = 100. The steady ROC computed from averaging 5,000 inferences is presented in a solid black line. The naive classifier is in cyan. The yellow box summarizes the statistics of the 50 ROC curves when compared to the steady ROC. Abbreviations: MC - Monte Carlo, ROC - receiver operating characteristic, AUC - area under the curve, and SSE - sum of the squared errors.

**Fig. 14.**
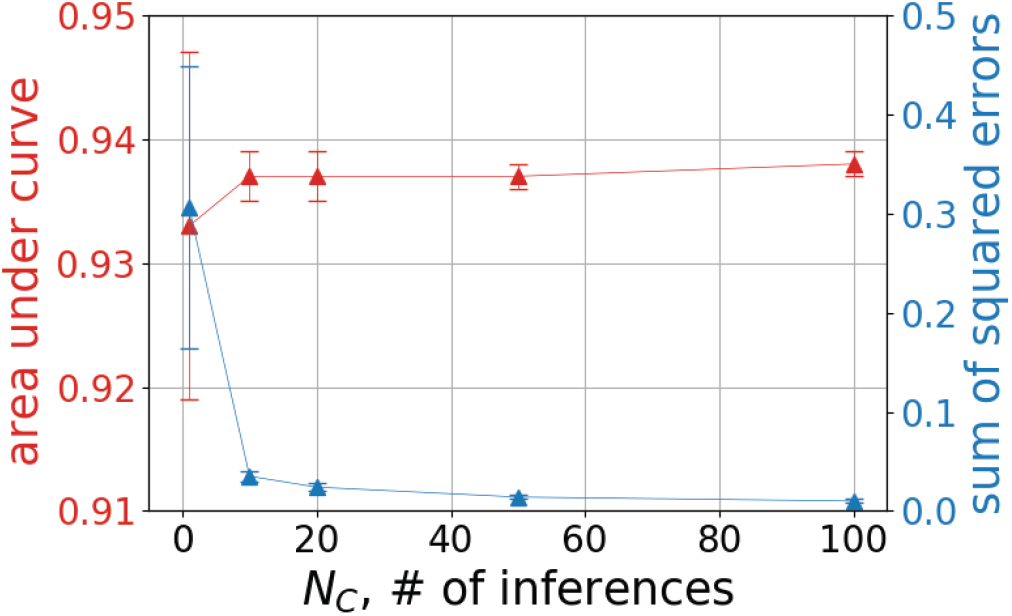
Summary statistics for candidate number of inferences. We computed AUC and sum of squared errors from 50 ROC curves generated by varying *N*_*C*_, the number of candidate MC dropout inferences. The AUC is plotted in red and the sum of squared errors is plotted in blue with corresponding color-coded vertical axes. Abbreviations: MC - Monte Carlo, ROC - receiver operating characteristic, and AUC - area under the curve.

**Fig. 15.**
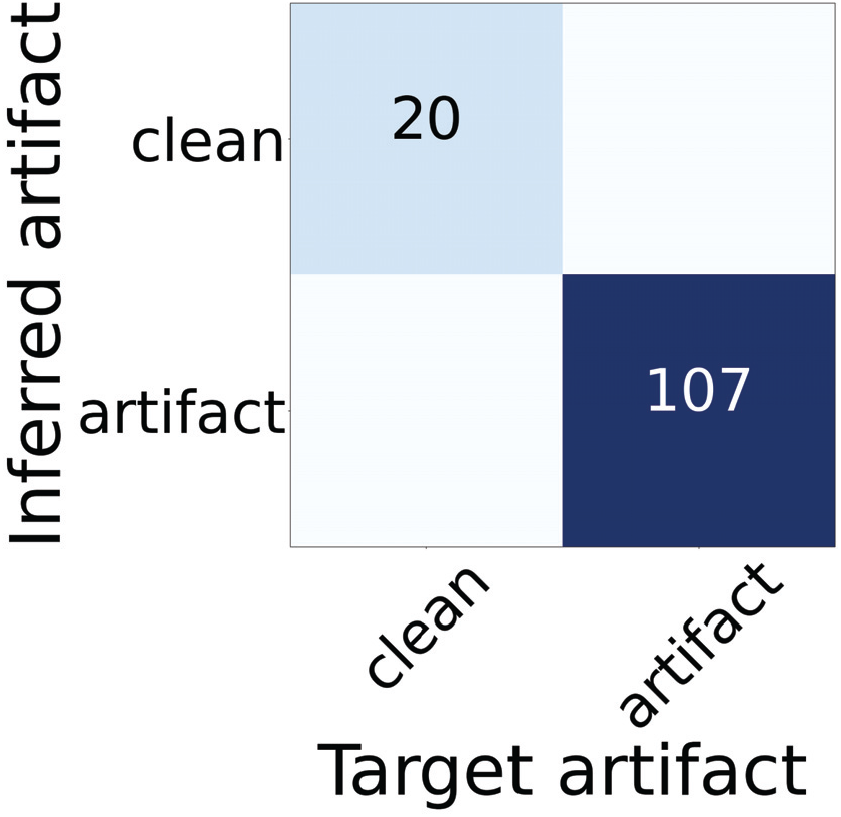
Stage I confusion matrix. The target was taken from the ground truth label obtained by visual inspection. The inferred artifact was determined using the proposed algorithm. All predictions appear along the diagonal, TP=107, TN=20, ACC = 1.0 and F1-score = 1.0.

**Fig. 16.**
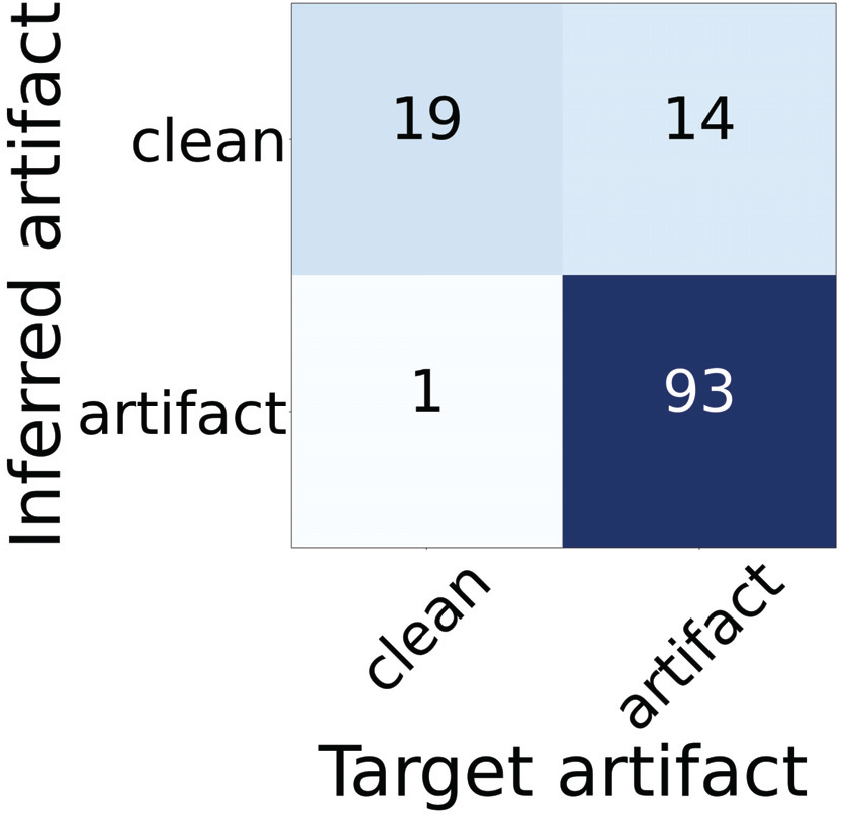
Stage I confusion matrix after applying ComBat (Fortin et al., 2018) to the proof-of-concept dataset. The target was taken from the ground truth label obtained by visual inspection. The inferred artifact was determined using the proposed algorithm. TP=93, FN=14, FP=1, TN=19, ACC = 0.882 and F1-score = 0.925.

**Fig. 17.**
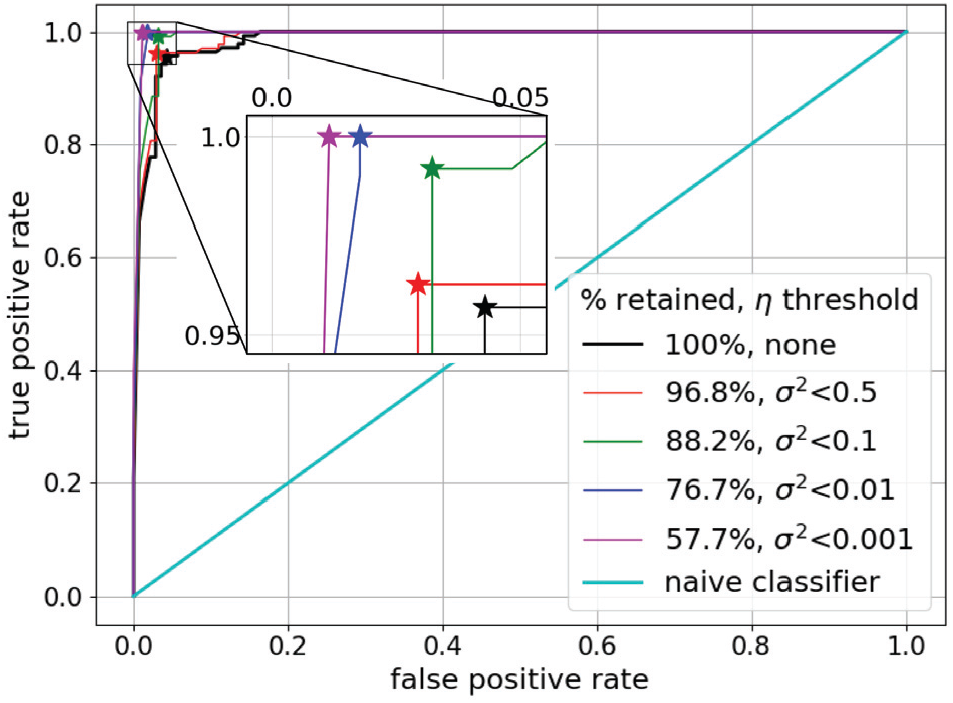
Stage II ROC plots with the implemented uncertainty. We computed 100 MC inferences on the validation subset to estimate *p* and σ^2^. We used σ^2^ < η to screen volumes of low confidence, i.e., high σ^2^, (and send them for visual inspection). The bottom-right inset specifies the percent retained after the corresponding η value is applied. The inflection point *J* is denoted with a color-coded star for each curve. The upper left inset is a zoomed-in version to highlight how *J* changes as η is lowered. Abbreviations: ROC - receiver operating characteristic, MC - Monte Carlo, *p* - prediction probability (Eq. 3), σ^2^ - sample variance (Eq. 5), η - uncertainty threshold, and *J* - inflection point (Eq. 4).

**Fig. 18.**
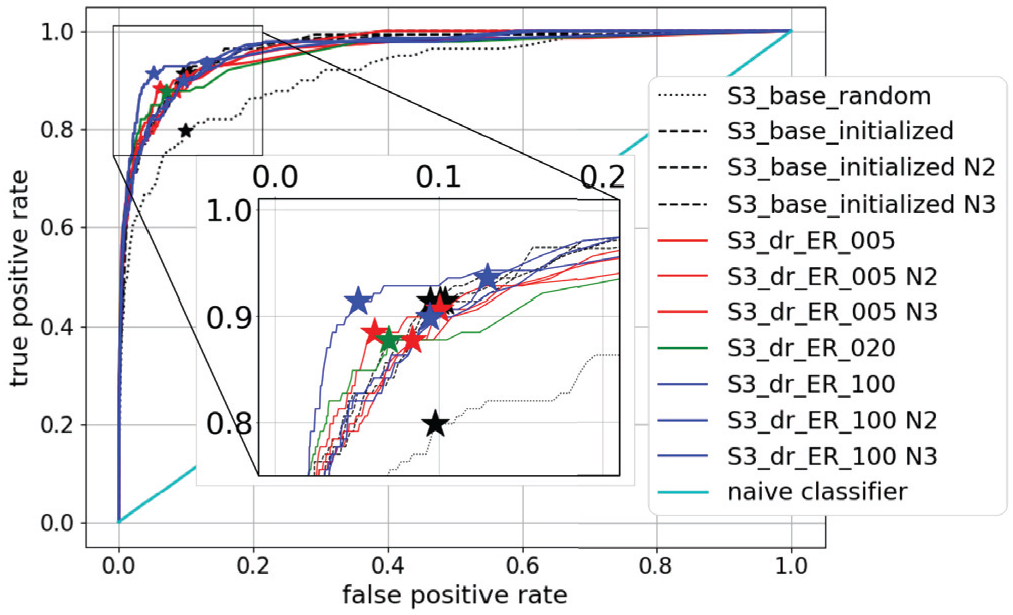
Stage III ROC plots for five comparison experiment types, including repeats. For each of the 11 comparison experiments, we computed 50 MC inferences on the validation subset. We computed *p* and set a classification threshold to generate ROC plots. We repeated three experiments – `S3_base_initialized`, `S3_dr_ER_005`, and `S3_dr_ER_100` – three times. The second and third repeats are denoted with N2 and N3. Abbreviations: ROC - receiver operating characteristic, MC - Monte Carlo, and *p* - prediction probability (Eq. 3).

**Fig. 19.**
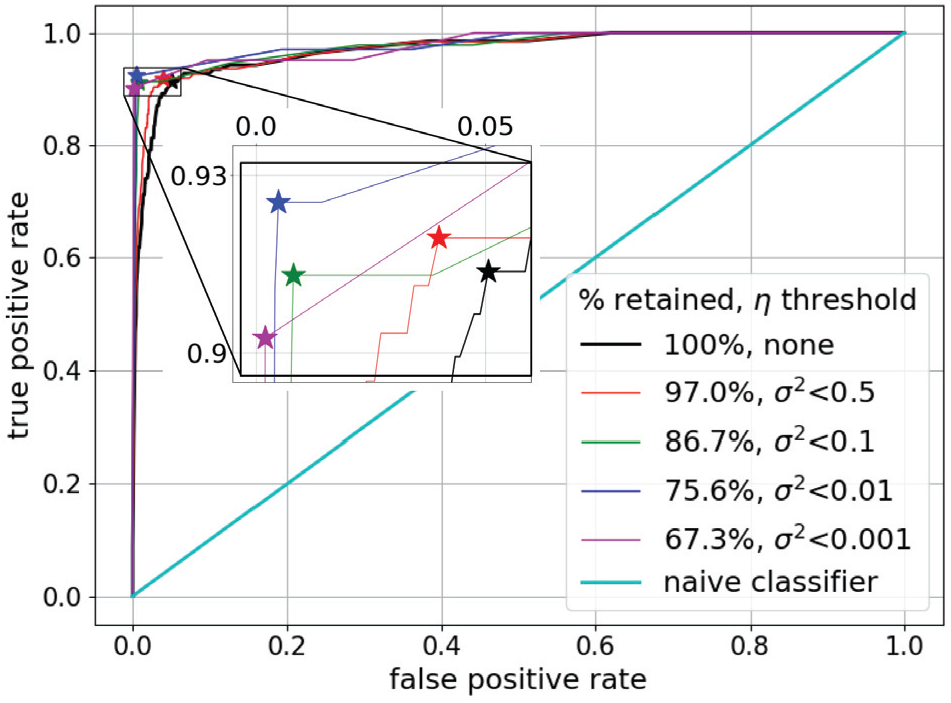
Stage III ROC plots with the implemented uncertainty. Following the `S3_dr_ER_100` experiment we computed 50 MC inferences on the validation subset to estimate prediction probability *p* and sample variance σ^2^. We set an uncertainty threshold η, on the normalized σ^2^, to screen volumes of low confidence, i.e., high σ^2^. The bottom-right inset specifies the percent retained after the corresponding η value is applied. The inflection point *J*, is denoted with a color-coded star for each curve. The upper-left inset is a zoomed-in version to highlight how the inflection point changes as η is lowered. Abbreviations: ROC - receiver operating characteristic, MC - Monte Carlo, *p* - prediction probability (Eq. 3), σ^2^ - sample variance (Eq. 5), η - uncertainty threshold, and *J* - inflection point (Eq. 4).

For comparison to the data-ramping experiments, we investigated using a focal loss, namely focal cross-entropy, instead of categorical cross-entropy. Focal loss has been developed by Lin et al. (2017) for dense object detection to deal with heavily imbalanced classes, i.e., the target object appears less frequently than the other objects in an image. We implemented Tensorflow’s Keras’ Binary Focal Cross-Entropy (API: tensorflow-website). We designed the `S3_focal_loss` experiment by implementing the following sets of parameters: γ = 2.0, applyclass-balancing=True, α = 0.02. We initialized with random parameters and trained 50 epochs with the realistic training subset (Table 3) comprising *T*_*r*_ = 20880 MRI volumes (CP = 98%). Fig. 12 presents the loss (focal cross-entropy) and accuracy, plotted for the training and validation subsets. After training completed, the model parameters were used to predict the data from corresponding validation and testing subsets, with results presented as a confusion matrix (Fig. 20).

**Fig. 20.**
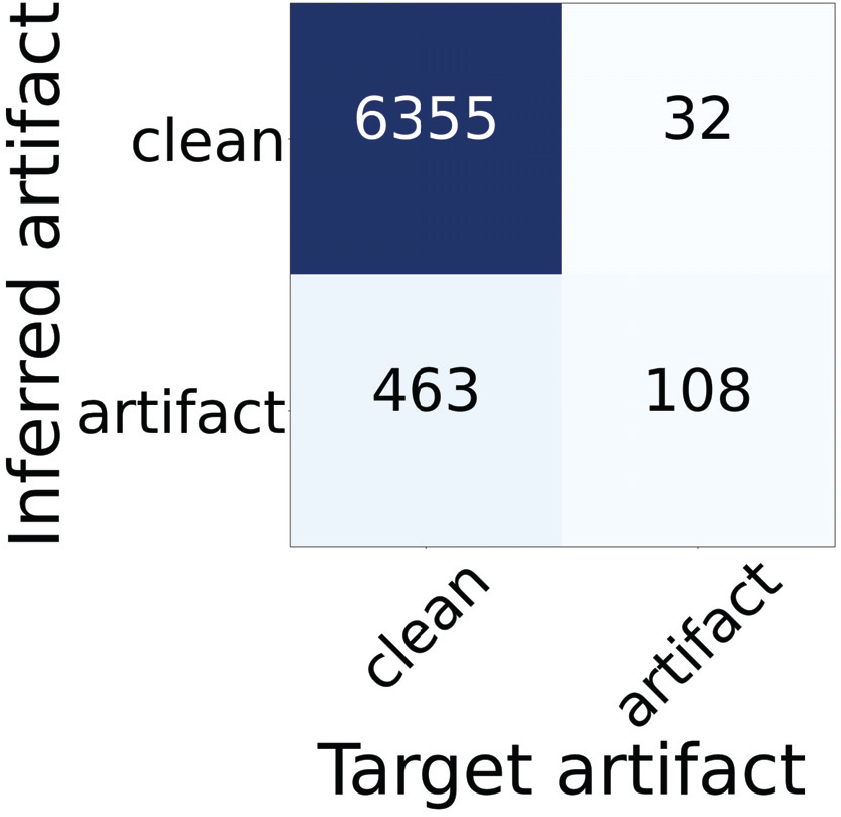
Stage III confusion matrix after using focal loss (Lin et al., 2017) to train with the realistic dataset. The target was taken from the ground truth label obtained by visual inspection. The inferred artifact was determined using the proposed algorithm after training with focal loss. TP=108, FN=32, FP=463, TN=6335, ACC = 0.929 and F1-score = 0.304.

### 2.4. Algorithm performance evaluation

After the training completed, we evaluated the performance of the algorithm, as follows. In each of the three developmental stages, we used the trained model parameters to make inferences *Ŷ*, on MRI volumes of both the validation and testing subset. We processed the inferences *Ŷ*, to classify the MRI volumes as either clean or artifact. In stage I, we computed the commonly used argmax of the output. In stages II and III, we developed a threshold approach detailed in Section 2.4.1, based on the probability and uncertainty generated by implementing MC dropout. After processing the inferences, we compared the binary classification to the ground truth *Y*. For comparing to the ground truth *Y*, we summarized the results by implementing the metrics commonly used for evaluating the diagnostic capacity of a binary classifier. That is, we developed the algorithm for artifact detection; we therefore defined positive detection to be the artifact class and negative detection to be the clean class, i.e., absence of artifact. These definitions then allowed us to set up the following terms summarized in Table 5 to guide the performance evaluation.

**Table 5.**
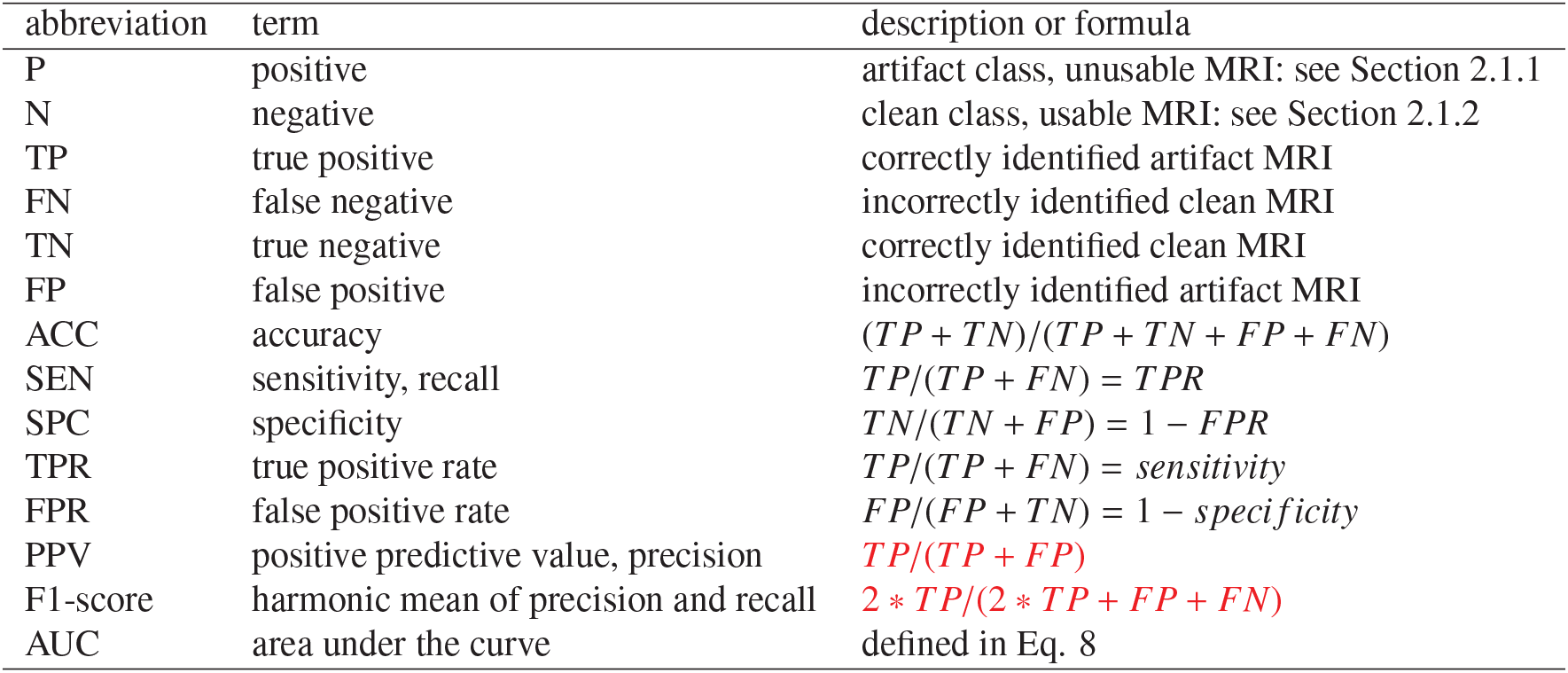
Summary of terms used for algorithm performance evaluation.

**Table 6.** Stage II summary statistics on the validation and testing subsets computed using inflection point *J*, for several values of the uncertainty threshold η, on the sample variance σ^2^. A lower uncertainty threshold is a stricter threshold. Abbreviations: *J* - inflection point (Eq. 4), η - uncertainty threshold, σ^2^ - sample variance (Eq. 5), AUC - area under the curve (Eq. 8), F1 - harmonic mean of precision and recall, ACC - accuracy, SPC - specificity, TN - true negative, FP - false positive, SEN - sensitivity, FN - false negative, TP - true positive, negative - clean MRI volume (usable), and positive - artifact MRI volume (unusable).

| set | $\eta$ | % retained | AUC | F1 | $J$ | ACC | SPC | TN | FP | SEN | FN | TP |
| --- | --- | --- | --- | --- | --- | --- | --- | --- | --- | --- | --- | --- |
| validation | none | 100.0 | 0.988 | 0.957 | 0.58 | 0.957 | 0.957 | 134 | 6 | 0.957 | 6 | 133 |
|  | 0.5 | 96.8 | 0.990 | 0.966 | 0.69 | 0.967 | 0.971 | 132 | 4 | 0.963 | 5 | 129 |
|  | 0.1 | 88.2 | 0.995 | 0.980 | 0.94 | 0.980 | 0.968 | 120 | 4 | 0.992 | 1 | 121 |
|  | 0.01 | 76.7 | 0.999 | 0.990 | 0.97 | 0.991 | 0.982 | 111 | 2 | 1.000 | 0 | 101 |
|  | 0.001 | 57.7 | 1.000 | 0.993 | 0.99 | 0.994 | 0.989 | 86 | 1 | 1.000 | 0 | 74 |
| testing | none | 100.0 | N/A | 0.912 | 0.58 | 0.910 | 0.899 | 125 | 14 | 0.921 | 11 | 129 |
|  | 0.01 | 71.0 | N/A | 0.980 | 0.97 | 0.980 | 0.989 | 94 | 1 | 0.971 | 3 | 100 |

**Table 7.** Stage III summary statistics on the validation and testing subsets computed using inflection point *J* for several values of the epistemic uncertainty threshold η on the sample variance σ^2^. A lower uncertainty threshold is a stricter threshold. Abbreviations: *J* - inflection point (Eq. 4), η - uncertainty threshold, σ^2^ - sample variance (Eq. 5), AUC - area under the curve (Eq. 8), F1 - harmonic mean of precision and recall, ACC - accuracy, SPC - specificity, TN - true negative, FP - false positive, FN - false negative, SEN - sensitivity, TP - true positive, negative - clean MRI volume (usable), and positive - artifact MRI volume (unusable).

| set | $\eta$ | % retained | AUC | F1 | $J$ | ACC | SPC | TN | FP | SEN | FN | TP |
| --- | --- | --- | --- | --- | --- | --- | --- | --- | --- | --- | --- | --- |
| validation | none | 100.0 | 0.974 | 0.416 | 0.74 | 0.949 | 0.949 | 6474 | 345 | 0.914 | 12 | 127 |
|  | 0.5 | 97.0 | 0.977 | 0.454 | 0.83 | 0.959 | 0.960 | 6363 | 264 | 0.919 | 10 | 114 |
|  | 0.1 | 86.7 | 0.983 | 0.750 | 0.12 | 0.991 | 0.992 | 5890 | 48 | 0.913 | 8 | 84 |
|  | 0.01 | 75.6 | 0.988 | 0.805 | 0.97 | 0.994 | 0.995 | 5165 | 25 | 0.925 | 5 | 62 |
|  | 0.001 | 67.3 | 0.986 | 0.851 | 0.99 | 0.997 | 0.998 | 4636 | 9 | 0.902 | 4 | 37 |
| testing | none | 100.0 | N/A | 0.357 | 0.74 | 0.941 | 0.944 | 6435 | 384 | 0.814 | 26 | 114 |
|  | 0.01 | 72.5 | N/A | 0.834 | 0.97 | 0.995 | 0.997 | 4957 | 14 | 0.851 | 11 | 63 |

#### 2.4.1. Monte Carlo dropout to generate epistemic uncertainty measures

Epistemic uncertainty is generated when noise is introduced to the model architecture when making an inference on an input image (Shamsi et al., 2021). In stages II and III, we implemented MC dropout to estimate probability and epistemic uncertainties. DL tools for regression and classification do not capture probability, and a model prediction can be uncertain even with a high softmax output. Gal and Ghahramani (2016) developed MC dropout by showing that dropout in neural networks can be interpreted as a Bayesian approximation of a Gaussian process, a well-known probabilistic model. This means that for a single MRI volume *X*, we can use multiple (*N*) dropouts to sample *N* times from a network’s output for generating multiple inferences *Ŷ*= {*ŷ*_1_,…, *ŷ*_*N*_}, and approximate probability and uncertainties (Nair et al., 2020). We incorporated MC dropout into the architecture (Fig. 2) to compute the expectation value of *Ŷ* and estimate *p*, the prediction probability:

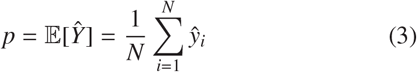

We set a classification threshold *p* > *t*, by varying over the range *t*∈[0, 1], to classify each MRI volume as positive (contains artifact) or negative (no artifact). We compared the classification to the ground truth *Y*, to compute TPR, FPR, and characterize the algorithm’s diagnostic capacity with ROC curves. We then computed *J*, the inflection point, as in (Pizarro et al., 2019):

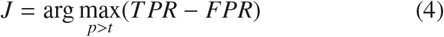

The inflection point is the (FPR, TPR) pair along the ROC curve that is closest to the optimal operating point, the top left corner of the plot. In addition to computing the probability, for each (*X,Ŷ*) pair, we computed three epistemic uncertainty metrics *u*_*m*_, with *m* = 1, 2, 3, following the methods in (Nair et al., 2020). The first uncertainty metric, the MC sample variance σ^2^, was estimated from the variance of the *N* inferences of the classification:

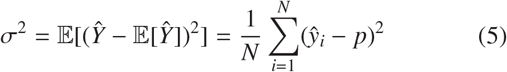

The second uncertainty metric, the predictive entropy termed *H*, was approximated across *N* inferences, and two possible labeled categories, *l* ∈ {0, 1}:

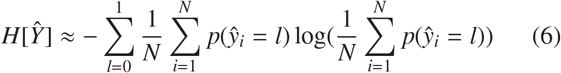

Finally, the third uncertainty metric, mutual information (*MI*), was approximated as the difference between *H*[*Ŷ*] and the expectation of the prediction entropies across inferences:

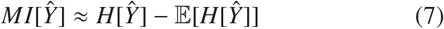

We rescaled each uncertainty metric, *u*_*m*_, to range between [0,1] by subtracting the minimum value in the subset and dividing by the range. We then set a pre-screen threshold, *u*_*m*_ < η, by varying over η = {0.5, 0.1, 0.01, 0.001}. We used this threshold to decide which MRI volumes to screen, i.e., send for visual inspection. The screened MRI scans were the ones that generated predictions with high uncertainty. We computed uncertainty *u*_*m*_, using the validation and testing subsets of the balanced dataset in stage II, and realistic dataset in stage III. For each uncertainty *u*_*m*_, decreasing the threshold η, resulted in more MRI scans sent out for visual inspection.

Epistemic uncertainty is generated when noise is introduced to the model architecture when making an inference on an input image (Shamsi et al., 2021). The complement of epistemic uncertainty is aleatoric uncertainty, generated when noise is introduced to the input image before making an inference with a static (unchanged) model. The model we implemented relies on epistemic uncertainty and MC dropout. However, implementing MC dropout to calculate epistemic uncertainty cannot be implemented by all DNN architectures. To demonstrate that our method can also apply to a variety of DNN architectures, we estimated the aleatoric uncertainty (Kendall and Gal, 2017). We considered using image processing techniques to introduce noise to the input image data to generate aleatoric uncertainty and the fact that certain image augmentations that can be used for this purpose could introduce MRI artifacts, potentially confusing the classifier. With this in mind, we opted for a conservative method, namely flipping the image along its three axes. We implemented flipping as a way to demonstrate the proof-of-concept for how to estimate aleatoric uncertainty. We introduced noise to the MRI data by flipping the image along its three axes. Flipping the image generates different images without altering the image intensity in any way. Importantly, it does not introduce image artifacts. Given the MRI volume has three axes (x,y,z), we toggled the flip: on or off. In total, each MRI volume generated 2^3^ = 8 variations of the same volume. We used the trained model for the best-performing experiment and removed the dropout component to generate inferences on eight variations for MRI volumes of the test set.

#### 2.4.2. Estimating the number of Monte Carlo inferences to characterize the performance

To accurately characterize performance with ROC curves generated by MC inferences, we estimated the number of inferences needed by exploring candidate values, *N*_*C*_ = {1, 10, 20, 50, 100}. To that end, we estimated a total of *N*_*T*_ = 1e4 inferences from the validation subset and split them in two. We used one half to compute a steady ROC curve defined as a stable ROC curve derived by implementing the law of large numbers, where the ROC curve gets closer to the expected value as the number of samples gets larger. To estimate the steady ROC, we first computed the steady probability *p*_*st*_, by plugging number of samples *N* = *N*_*T*_ /2 in Eq. 3. We set a classification threshold *p*_*st*_ > *t* to generate steady TPR and steady FPR to plot the steady ROC curve, shown in solid black line in Fig. 13. We then used the other half, *N*_*T*_ /2 inferences, to randomly sample 50 sets made up of *N*_*C*_ inferences. For each set, *j* = {1,…, 50}, we computed set probability *p*_*j*_, using number of samples *N* = *N*_*C*_ in Eq. (2). We set a classification threshold *p*_*j*_ > *t* to generate set TPR and set FPR to plot 50 ROC curves from each set, as presented in Fig. 13 for candidate values *N*_*C*_ = {1, 10, 100}. With increasing *N*_*C*_, the ROC curves in Fig. 13 converged more closely around the steady ROC as *N* increased, quantified for each set *j* with the following statistics. We approximated the set area under the curve *AUC*_*j*_, by integrating over classification threshold *t* ∈ [0, 1]:

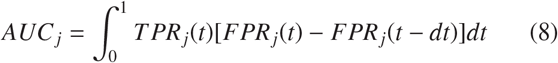

Next, we computed the set sum of the squared errors, *SS E* _*j*_ by integrating the squared error between each set *j*, and the steady estimate *st*, over classification threshold *t* ∈ [0, 1]:

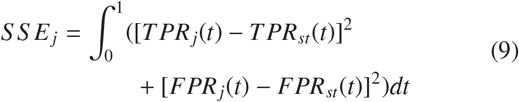

Fig. 14 presents the estimated mean (denoted with triangles) and standard deviation (denoted with vertical bars) for *AUC*_*j*_ and *SS E* _*j*_ over 50 sets, as a function of the candidate values, *N*_*C*_. The changes in the *AUC* mean across the candidate values *N*_*C*_, were small; however, the variance decreased by a factor of approximately 10. The *SS E* mean decreased by a factor of approximately 10 between *N*_*C*_ = 1 and *N*_*C*_ = 10 and decreased further by a factor of approximately 3 between *N*_*C*_ = 10 and *N*_*C*_ = 100. To accurately characterize the performance, we used *N*_*C*_ > 50.

#### 2.4.3. Evaluating performance with bottom-line metrics

Investigators routinely report artifact detection results with performance metrics that can be misleading, such as those used here: AUC, accuracy, specificity, and sensitivity. Each metric provides an insight into classification performance; however, their relevance depends on the dataset size and the categorical balance (e.g. clean percentage). For example, classifying all MRI volumes as clean in a 98% clean dataset, would result in an impressive 98% accuracy, with erroneous classification of all artifact MRI volumes. These routinely used performance metrics, although intuitive, can be misleading without consideration for the larger context. In our study, by implementing uncertainties our results improved at the cost of cases sent for visual inspection and generating additional manual work, a metric not used in the literature. In addition to reporting the commonly used metrics, Alfaro-Almagro et al. (2018) evaluated their method by including two metrics that are specific to artifact detection and provide a bottom-line to the performance of the algorithm: unusable datasets missed (UDM) and datasets flagged for manual review (DFFMR). UDM estimates how ‘dirty’ the dataset will remain upon implementation of the algorithm, as inevitably some MRI volumes with artifacts will go undetected. DFFMR is the percentage of MRI volumes that were classified as potential artifacts – i.e. flagged for inspection – estimating the amount of manual work needed to reach UDM. Alfaro-Almagro et al. (2018) selected a threshold such that they would review 17.29% of MRI volumes (DFFMR), resulting in missing 0.15% of MRI volumes with artifacts (UDM).

## 3. Results

We developed a DL algorithm to automatically identify the artifact of a brain MRI. In stage I, we summarized the prediction with a confusion matrix, to illustrate the performance on a small dataset. In stage II, we used ROC plots to illustrate how uncertainty metrics can improve the prediction performance; then, we estimated testing errors. In stage III, we compared the five types of experiments using ROC plots, incorporated UDM versus DFFMR, and estimated test errors for the winning experiment.

### 3.1. Stage I: Testing the proof-of-concept with a small dataset

In stage I, we identified a dataset to test the proof-of-concept with the proposed DL algorithm. After training, we used the model to infer on each MRI volume from the testing subset and compared to the ground truth. We generated a confusion matrix (Fig. 15) to visualize the performance, as in (Pizarro et al., 2019). The DL algorithm matched the ground truth on all MRI volumes with accuracy = 1.0 (F1-score = 1.0, specificity = 1.0 and sensitivity = 1.0). Although the dataset was small and beset by issues addressed in the latter stages, it was encouraging to see DL distinguish clean MRI volumes from artifact MRI volumes. To address the issue of domain difference across clean MRI and artifact MRI, we applied Combat to the proof-of-concept dataset. After training, we used the model to infer on each MRI volume from the testing subset and compared to the ground truth. We generated a confusion matrix (Fig. 16) to visualize the performance. The performance is summarized with accuracy = 0.882 (F1-score = 0.925, specificity = 0.950 and sensitivity = 0.869). The performance decreased slightly relative to the results obtained before domain adaptation, although the results remained encouraging.

### 3.2. Stage II: Making the dataset balanced and developing uncertainties

We built on the encouraging results obtained in stage I. After a few unsuccessful training sessions with the realistic dataset, we created stage II to continue developing our algorithm. We summarized the results as follows. To implement MC dropout for generating probabilities and uncertainties we created a balanced dataset with CP=50%. We saved the model parameters after the stage II training completed, then used the model to make *N* = 100 inferences on the validation and testing subsets. We used the *N* inferences to compute prediction probability *p* (Eq. 3) and three uncertainties *u*_*m*_: sample variance σ^2^, entropy *H*, and mutual information *MI*, following Eqs. 5-7. We rescaled the uncertainties and set a pre-screen threshold *u*_*m*_ < η, by varying over η = {0.5, 0.1, 0.01, 0.001}. The prescreen threshold was used to determine which MRI volumes to send for visual inspection. MRI volumes sent for visual inspection had high uncertainty, i.e., the prediction confidence was low relative to the confidence attached to the other volumes in the subset. For each η we used *p* > *t* to generate five ROC curves, without and with pre-screen threshold, as illustrated for the sample variance σ^2^ < η in Fig. 17. To evaluate the change in performance as the pre-screen threshold η, was lowered (meaning the threshold became stricter), we computed the inflection point *J*, following Eq. 4, and summarized in Table 6. As η was lowered, the percent retained decreased (more volumes were sent for visual inspection), the AUC increased, and *J* moved closer to the top left, with increased classification confidence. We used the validation subset results to select (η, *J*) = [(*none*, 0.58), (0.01, 0.97)] and classify the testing subset with and without a pre-screen threshold (included in bottom two rows of Table 6).

### 3.3. Stage III: Implementing transfer learning for classifying a realistic – 98% clean – dataset

Implementing uncertainties in stage II allowed us to obtain high classification performance. In stage III, to transfer the learning from stage II and generate results on a realistic dataset, we extended the dataset to CP=98%. Our initial attempts to ramp the clean percentage resulted in limited performance, as shown in Fig. 8. We therefore designed five comparison experiment types focused on optimizing the ramping of the dataset size. We repeated three of the five comparison experiment types two additional times, for a total of: 5 + 3 * 2 = 11 experiments. After training the 11 comparison experiments (described in Section 2.3.3), we used each model to make *N* = 50 inferences on the validation subset. We used the *N* inferences to compute prediction probability *p* (Eq. 3), and set a classification threshold *p* > *t*, to generate 11 ROC plots, illustrated in Fig. 18. All the ROC curves in Fig. 15 appear to overlap, indicating similar levels of performance, except for the following two: `S3_base_random` with lower performance and `S3_dr_ER_100` with higher performance.

We selected `S3_dr_ER_100` as the winning model and used the *N* = 50 inferences to estimate sample variance σ^2^ (Eq. 5). We rescaled σ^2^ and set a pre-screen threshold σ^2^ < η, by varying over η = {0.5, 0.1, 0.01, 0.001}. For each η, we used *p* > *t* to generate five ROC curves, with and without pre-screen threshold, illustrated in Fig. 19. The pre-screen threshold was used to determine which MRI volumes to send for visual inspection, as the uncertainty was high, i.e., the prediction confidence was relatively lower than the confidence achieved for the other volumes in the subset. We computed the inflection point *J* (Eq. 4) and summarized in Table 7, to compare the performance obtained as the pre-screen threshold η was lowered (meaning the threshold became stricter). As η was lowered, the percent retained decreased (more volumes were sent for visual inspection), the AUC increased, and the inflection point *J* moved in the general direction to the left. The AUC for the curves in Fig. 19 was lower than those in Fig. 17, as the class imbalance increased, causing the ROC curves to be more skewed to lower true positive rate values, i.e., lower sensitivity. We used the saved `S3_dr_ER_100` model to make *N* = 100 inferences on the testing subset and estimate prediction probability *p* and sample variance σ^2^. We used the validation subset results to select (η, *J*) = [(*none*, 0.74), (0.01, 0.97)] and classify the testing subset with and without a pre-screen threshold (included in bottom two rows of Table 7).

To compare our data-ramping technique for handling class imbalance, we investigated training the algorithm with focal loss (Lin et al., 2017) that was developed for a similar purpose. After training with the realistic, we used the model to infer on each MRI volume from the testing subset and compared to the ground truth. We generated a confusion matrix (Fig. 20) to visualize the performance. The performance is summarized with accuracy = 0.929 (F1-score = 0.304, specificity = 0.932 and sensitivity = 0.771) and included in Table 9 for comparison.

**Table 8.**
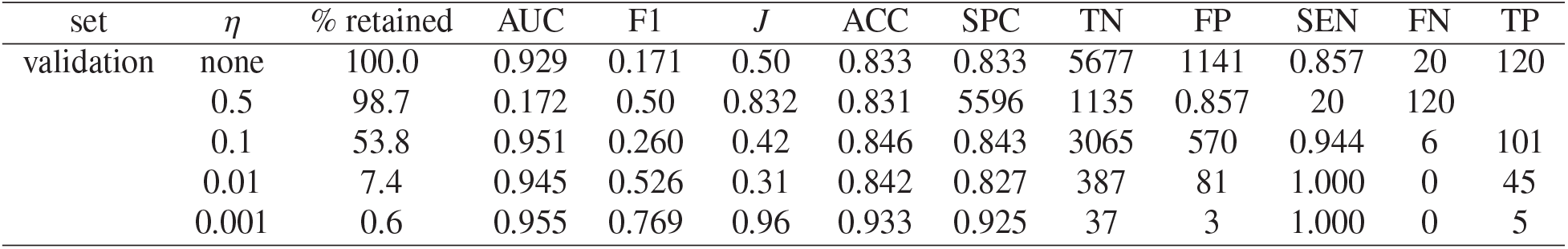
Stage III summary statistics on the validation subsets computed using inflection point *J* for several values of the aleatoric uncertainty threshold η on the sample variance σ^2^. A lower aleatoric uncertainty threshold is a stricter threshold. Abbreviations: *J* - inflection point (Eq. 4), η - uncertainty threshold, σ^2^ - sample variance (Eq. 5), AUC - area under the curve (Eq. 8), F1 - harmonic mean of precision and recall, ACC - accuracy, SPC - specificity, TN - true negative, FP - false positive, FN - false negative, SEN - sensitivity, TP - true positive, negative - clean MRI volume (usable), and positive - artifact MRI volume (unusable).

| set | $\eta$ | % retained | AUC | F1 | $J$ | ACC | SPC | TN | FP | SEN | FN | TP |
| --- | --- | --- | --- | --- | --- | --- | --- | --- | --- | --- | --- | --- |
| validation | none | 100.0 | 0.929 | 0.171 | 0.50 | 0.833 | 0.833 | 5677 | 1141 | 0.857 | 20 | 120 |
|  | 0.5 | 98.7 | 0.172 | 0.50 | 0.832 | 0.831 | 5596 | 1135 | 0.857 | 20 | 120 |  |
|  | 0.1 | 53.8 | 0.951 | 0.260 | 0.42 | 0.846 | 0.843 | 3065 | 570 | 0.944 | 6 | 101 |
|  | 0.01 | 7.4 | 0.945 | 0.526 | 0.31 | 0.842 | 0.827 | 387 | 81 | 1.000 | 0 | 45 |
|  | 0.001 | 0.6 | 0.955 | 0.769 | 0.96 | 0.933 | 0.925 | 37 | 3 | 1.000 | 0 | 5 |

**Table 9.** Table of ablation to compare how performance on the validation set was impacted by various aspects incorporated in the algorithm. Abbreviations: CCE - categorical cross-entropy, FCE - focal cross-entropy, LR - learning rate, CLR - cyclical learning rate, CW - class weights, TL - transfer learning, DR - data ramping, MC - monte carlo

| Experiment<br>(aspect introduced) | LR schedule |  | Handling class imbalance |  |  |  | Uncertainty |  |
| --- | --- | --- | --- | --- | --- | --- | --- | --- |
|  | LR | CLR | CW | TL | DR | FCE | epistemic | aleatoric |
| CCE | X | X | X | X | X |  | X | X |
| FCE |  |  |  |  |  | X |  |  |
| linear LR | X |  |  |  |  | X |  |  |
| CLR |  | X | X | X | X |  | X | X |
| CW |  |  | X | X | X |  | X | X |
| TL |  |  |  | X | X |  | X | X |
| DR |  |  |  |  | X |  | X | X |
| epistemic |  |  |  |  |  |  | X |  |
| aleatoric |  |  |  |  |  |  |  | X |
| accuracy | 0.893 | 0.989 | 0.900 | 0.897 | 0.949 | 0.929 | **0.994** | 0.846 |
| precision | 0.134 | 0.859 | 0.143 | 0.152 | 0.269 | 0.189 | **0.713** | 0.151 |
| recall | 0.799 | 0.519 | 0.799 | 0.914 | 0.914 | 0.771 | 0.925 | **0.944** |
| F1 | 0.230 | 0.647 | 0.242 | 0.261 | 0.416 | 0.304 | **0.805** | 0.260 |
| AUC | 0.926 | 0.932 | 0.921 | 0.968 | 0.974 | 0.964 | **0.988** | 0.951 |

**Table A.10.** Clean Selector: two-step process to select the number of MRI volumes required in stage II to maintain the distribution of the artifact dataset and study-wise 50% clean. Abbreviations: **s** - study number, **v**_*a*_ - # of MRI with artifact, 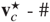 of clean MRI needed to ensure study-wise 50% clean, 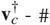 of clean MRI available, **v**_*R*_ - 137 missing MRI volumes redistributed to keep studywise 50% clean, as explained in text, **v**_*c*_ - number of clean MRI volumes taken, and **cp** - final clean percent.

| $s$ | $v_a$ | $v_c^*$ | $v_c^\dagger$ | $v_R$ | $v_c$ | $cp$ |
| --- | --- | --- | --- | --- | --- | --- |
| 1 | 141 | 141 | 18481 | 36 | 177 | 55.7% |
| 2 | 91 | 91 | 20366 | 22 | 113 | 55.4% |
| 3 | 90 | 90 | 0 | 0 | 0 | 0.0% |
| 4 | 89 | 89 | 8135 | 22 | 111 | 55.5% |
| 5 | 83 | 83 | 23881 | 20 | 103 | 55.4% |
| 6 | 38 | 38 | 0 | 0 | 0 | 0.0% |
| 7 | 37 | 37 | 6905 | 9 | 46 | 55.4% |
| 8 | 29 | 29 | 5991 | 7 | 36 | 55.4% |
| 9 | 27 | 27 | 3956 | 7 | 34 | 55.7% |
| 10 | 13 | 13 | 883 | 3 | 16 | 55.2% |
| 11 | 13 | 13 | 742 | 3 | 16 | 55.2% |
| 12 | 11 | 11 | 489 | 3 | 14 | 56.0% |
| 13 | 7 | 7 | 5713 | 2 | 9 | 56.3% |
| 14 | 6 | 6 | 1222 | 1 | 7 | 53.9% |
| 15 | 6 | 6 | 1441 | 1 | 7 | 53.9% |
| 16 | 5 | 5 | 0 | 0 | 0 | 0.0% |
| 17 | 4 | 4 | 776 | 1 | 5 | 55.6% |
| 18 | 2 | 2 | 0 | 0 | 0 | 0.0% |
| 19 | 2 | 2 | 2229 | 0 | 2 | 50.0% |
| 20 | 2 | 2 | 0 | 0 | 0 | 0.0% |
| **Total** | **696** | **696** | **101210** | **137** | **696** | **50.0%** |

We summarized the results of the inferences with aleatoric uncertainties as ROC curves, presented in Fig. 21 in two ways. On the top, we isolated the prediction based on the orientation of the MRI volume, where: 000 indicates the original image, 001 indicates a flip along the z-axis, 010 a flip along the y-axis, 100 a flip along the x-axis, and so forth, until 111 where we flipped all directions x, y, and z. The algorithm is tolerant of the flips and the performance varies minimally for each orientation presented. On the bottom, we summarized the prediction in a similar way as with epistemic uncertainty. First, we computed a probability as the average of all eight predictions. Next, we computed a normalized sample variance, σ^2^, and used a threshold to remove uncertain volumes. We computed the inflection point *J* (Eq. 4) and summarized in Table 8, to compare the performance obtained as the pre-screen threshold η was lowered (meaning the threshold became stricter). As η was lowered, the percent retained decreased (more volumes were sent for visual inspection), the AUC increased, and the inflection point *J* moved in the general direction to the top. Compared to Table 7, applying similar levels of threshold removed more of the MRI volumes. This could be attributed to summarizing over eight instances of the volume instead of 50. Regardless, these ROC plots demonstrate the capabilities of implementing aleatoric uncertainty to the proposed algorithm. We included σ^2^ < 0.1 results in Table 9 for comparison. Note: We omitted the unimpressive test results.

**Fig. 21.**
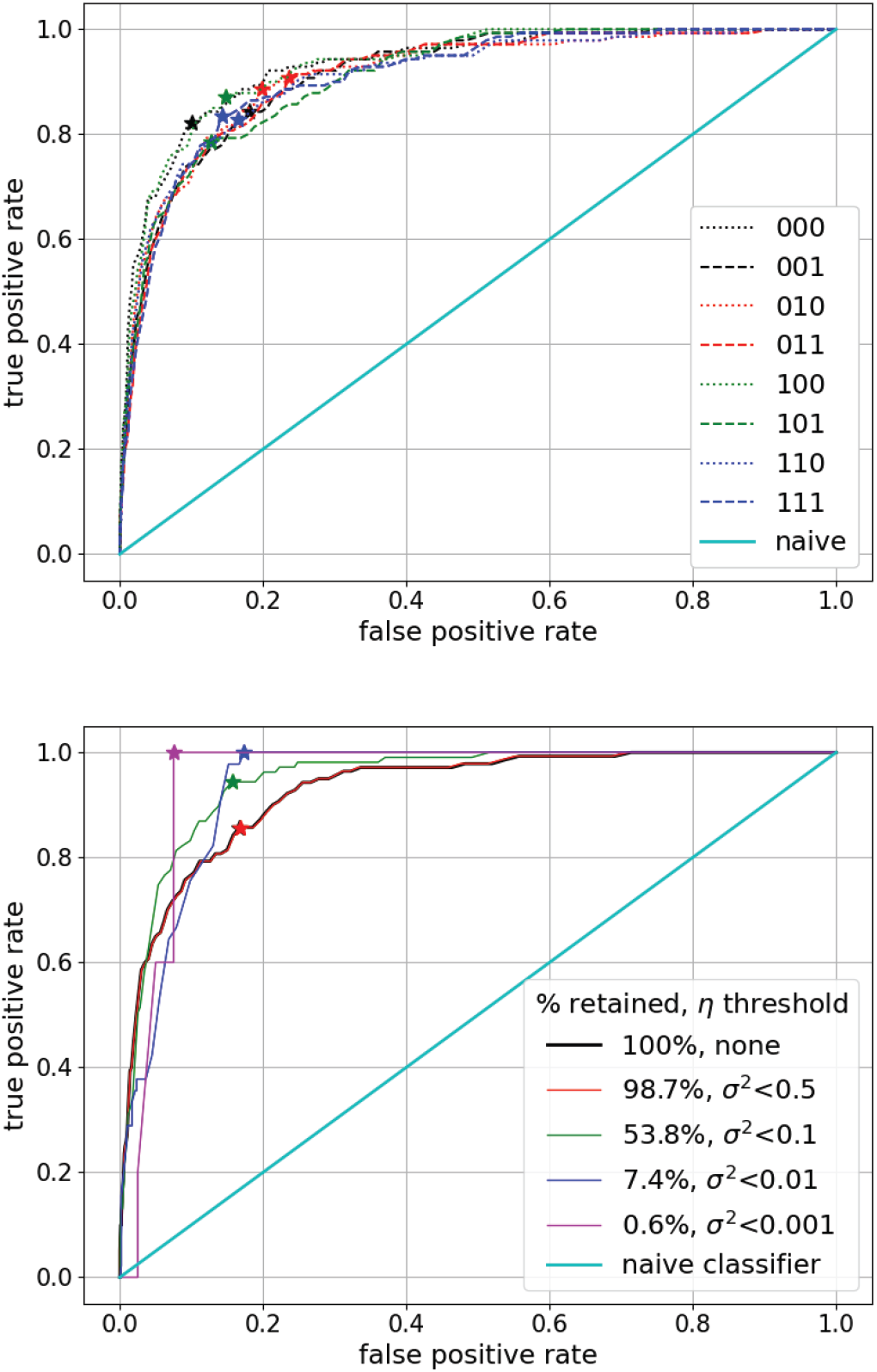
ROC with aleatoric uncertainties. On the top, we isolated the prediction based on the orientation of the MRI volume, where: 000 indicates the original image, 001 indicates a flip around the z-axis, 010 a flip around the y-axis, 100 a flip around the x-axis, and so forth, until 111 where we flipped around all axes x, y, and z. At the bottom, we summarized the prediction with a threshold on the sample variance similar to Fig. 19. We computed 8 aleatoric inferences on the validation subset to estimate prediction probability *p* and sample variance σ^2^. We set an aleatoric uncertainty threshold η, on the normalized σ^2^, to screen volumes of low confidence, i.e., high σ^2^. The bottom-right inset specifies the percent retained after the corresponding η value is applied. The inflection point *J* is denoted with a color-coded star for each curve. Abbreviations: ROC - receiver operating characteristic, *p* - prediction probability (Eq. 3), σ^2^ - sample variance (Eq. 5), η - uncertainty threshold, and *J* - inflection point (Eq. 4).

After obtaining high-performance results by implementing epistemic uncertainties in stages II and III, we wanted to compare our results to those obtained by Alfaro-Almagro et al. (2018), the current state-of-the-art in automated artifact detection. Fig. 22 presents curves of UDM versus DFFMR, two insightful bottom-line metrics for MRI artifact detection, obtained for 143 combinations as follows. We computed UDM and DFFMR for each of the 11 comparison experiments (including repeats, see Section 2.3.3 and Fig. 18 obtained with and without a pre-screen threshold. The pre-screen threshold *u*_*m*_ < η was applied for three uncertainty metrics (sample variance, entropy, and mutual information), and four threshold values η = {0.5, 0.1, 0.01, 0.001}. This resulted in each of the 11 experiments generating 1 + 3(4) = 13 combinations (without and with pre-screen threshold) represented by a marker in Fig. 22. Similar to the ROC curve, a tradeoff between UDM and DFFMR can be observed. As η was lowered (meaning the threshold became stricter), UDM decreased and DFFMR increased. Setting the result obtained by Alfaro-Almagro et al. (2018) as a reference, the majority of the points generated by our study are located below and/or to the left, in line with DL outperforming machine learning. We highlighted two points with a red square and circle in Fig. 22, for comparison to the results obtained by Alfaro-Almagro et al. (2018). The results depicted by the red square generated approximately 50% reduction in artifact detection errors (UDM = 0.07%) relative to the results obtained by Alfaro-Almagro et al. (2018), with approximately the same percentage of datasets sent for manual review (DFFMR = 17.28%). The results depicted by the red circle generated a similar level of artifact detection (UDM = 0.14%) with less than half the manual work (DFFMR = 8.41%).

**Fig. 22.**
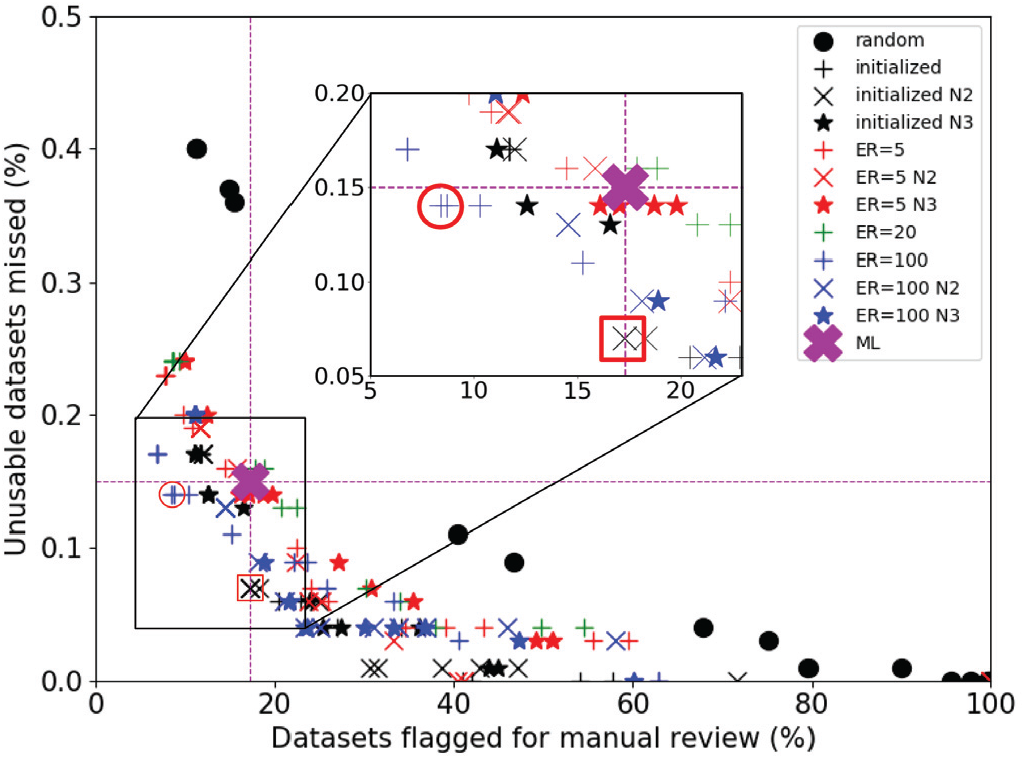
UDM versus DFFMR. The UDM and DFFRM were obtained from 143 combinations used in 11 experiments, three uncertainty metrics *u*_*m*_, and four threshold values η. Each experiment generated 13 combinations (with and without uncertainty threshold) indicated by a marker. Lowering the uncertainty threshold, η (meaning the threshold became stricter), caused the markers to move asymptotically down and to the right, indicating lower UDM at the expense of higher DFFMR. We selected two points – marked with a red circle and red square – to compare to the results obtained by ML (in purple X; (Alfaro-Almagro et al., 2018)). Abbreviations: UDM - unusable datasets missed, DFFMR - datasets flagged for manual review, and ML - machine learning.

To thoroughly lay out the aspects developed in this project, which have a positive impact on performance, we included a table of ablation (Liu et al., 2022; Aleef et al., 2021), in Table 9. We included the following aspects: LR schedule (CLR vs. linear LR), handling class imbalance (class weights, transfer learning, data ramping, focal loss), and uncertainty (epistemic and aleatoric). To compare these aspects and experiments, we included the following statistical metrics to summarize performance: accuracy, precision, recall, F1 score, and AUC. Regarding the two LR schedule experiments investigated, the performance of linear LR and CLR were comparable, with CLR generating slightly higher AUC than linear LR. In terms of handling class imbalance, class weights provided minimal increments to the performance. Transfer learning boosted the AUC and recall while maintaining similar levels in other metrics. Data ramping techniques outperformed all other methods developed for handling class imbalance. Focal loss outperformed class weights and generated higher accuracy, precision, and F1 when compared to transfer learning. Finally, implementing epistemic uncertainty, with MC dropout, proved the most impactful by generating extremely high performance.

## 4. Discussion

### 4.1. State-of-the-art comparison

We developed a DL algorithm to detect artifacts in a large, imbalanced brain MRI dataset. Our results are summarized as accuracy versus dataset size in Fig. 23, alongside other published results in brain MRI artifact detection. Our stage I results, depicted by a black dot, reproduced the results obtained using a small balanced dataset by (Küstner et al., 2018; Fantini et al., 2018; Oksuz, 2020); however, compared to these studies, we used a larger, imbalanced dataset. To generate large amounts of data, Küstner et al. (2018) and Fantini et al. (2018) split the MRI volumes into smaller patches, and Oksuz (2020) generated synthetic artifacts. In contrast, we used a large, imbalanced database of real MRI volumes. When working with a small, balanced dataset, visual inspection is more manageable and should be considered as the preferred method. The real challenge is implementing these automated methods in large and imbalanced neuroimaging databases. This was previously attempted in several studies (Esteban et al., 2017; Pizarro et al., 2016; Alfaro-Almagro et al., 2018; Mortamet et al., 2009; Sujit et al., 2019) whose performance is plotted in red in Fig. 23. In stage II, we used a balanced dataset of similar size and generated higher accuracies (plotted with left and right black triangles in Fig. 23). In stage III, we curated a realistic, imbalanced dataset that was larger than any previously reported work detecting artifact MRI, and we used it to generate 99.5% accuracy. We expect that the method in (Sujit et al., 2019) would benefit from larger datasets as DL architectures are large, composed of millions of parameters, and have been shown to improve in performance when more samples are presented (Pizarro et al., 2019). In contrast, the traditional machine learning approaches (Esteban et al., 2017; Pizarro et al., 2016; Alfaro-Almagro et al., 2018; Mortamet et al., 2009) may not benefit as much from larger datasets.

**Fig. 23.**
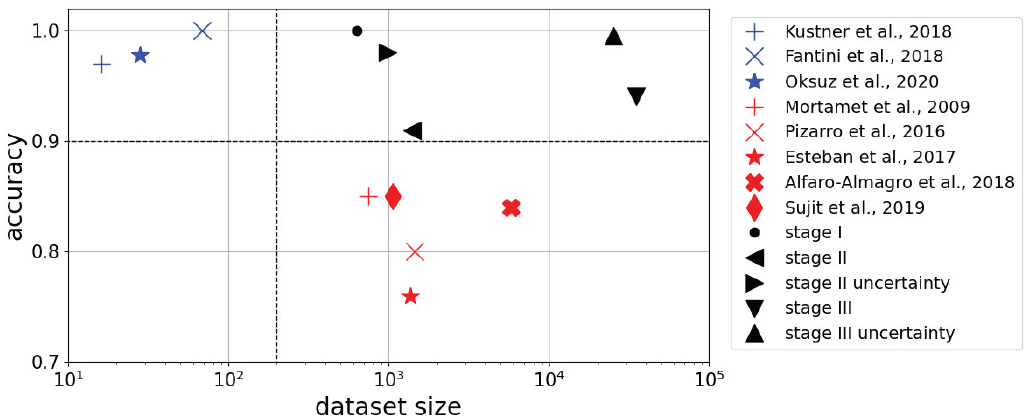
State-of-the-art in MRI artifact detection. Reported accuracy versus dataset size (log-scale) for studies that developed an automated algorithm. Studies in blue reported > 90% accuracy using small datasets (< 100 MRI volumes). Studies in red used larger datasets (> 700 MRI volumes) with accuracy < 90%. Our proposed algorithm generated accuracies > 90% over all three stages with datasets between 600 and 34,800 MRI volumes. Abbreviation: MRI - magnetic resonance imaging.

**Fig. B.24.**
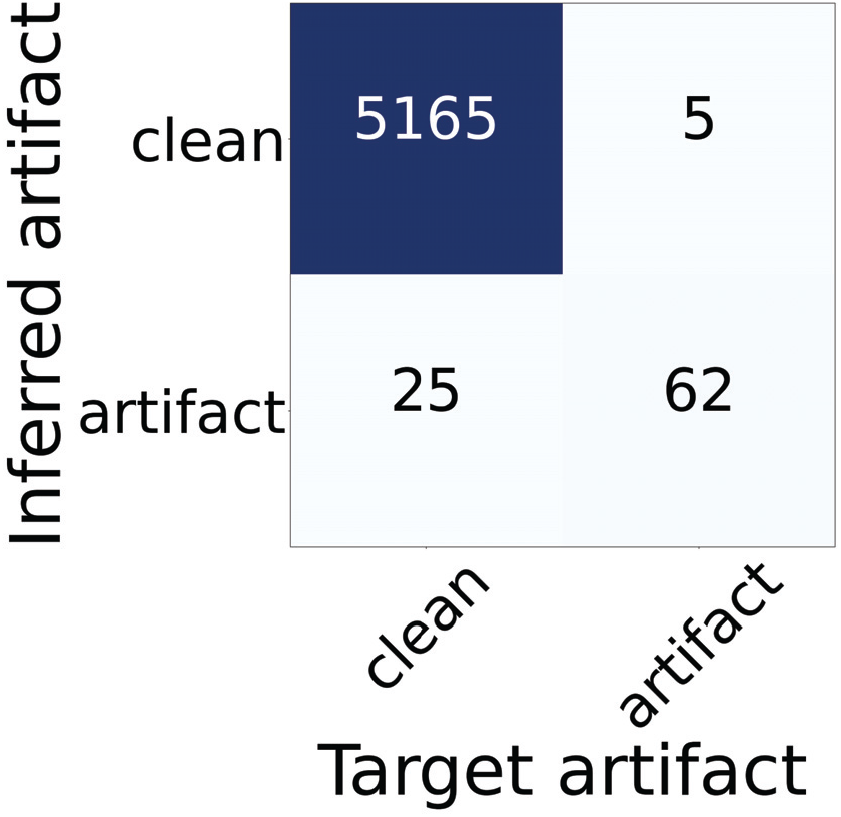
Stage III confusion matrix of the validation subset computed using inflection point *J* for several values of the epistemic uncertainty threshold η on the sample variance σ^2^ < 0.01. The predictions are summarized as F1 = 0.805, ACC = 0.994, SPC = 0.995, TN = 5165, FP = 25, SEN = 0.925, TP = 62, and FN = 5. Abbreviations: ACC - accuracy, SPC - specificity, TN - true negative, FP - false positive, SEN - sensitivity, FN - false negative, TP - true positive. Note that negative stands for clean MRI volume (usable), and positive stands for artifact MRI volume (unusable).

**Fig. B.25.**
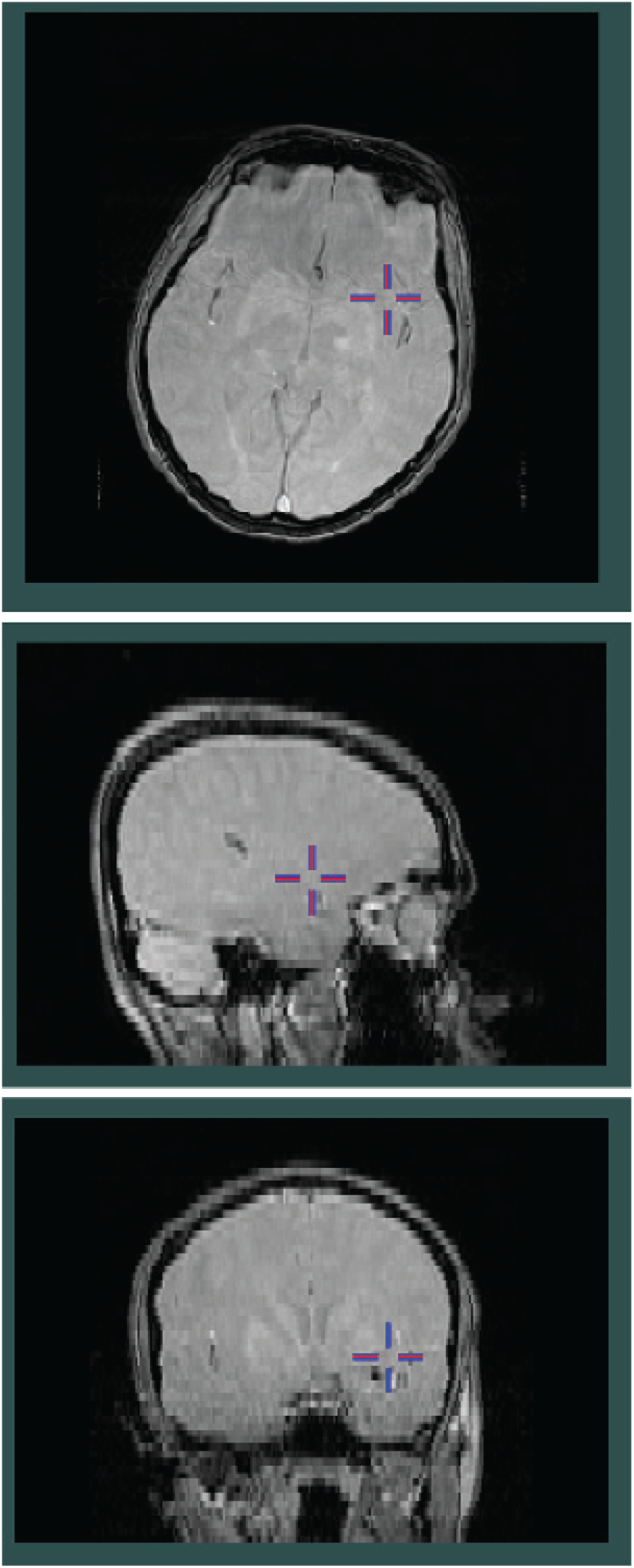
One of the two images out of the five that were previously labeled as artifacts. The image presents a magnetic transfer ON MRI contrast. As confirmed by an expert, this image is acceptable for analysis. The expert did not agree with the initial human rater’s label that the image showed a movement artifact.

**Fig. B.26.**
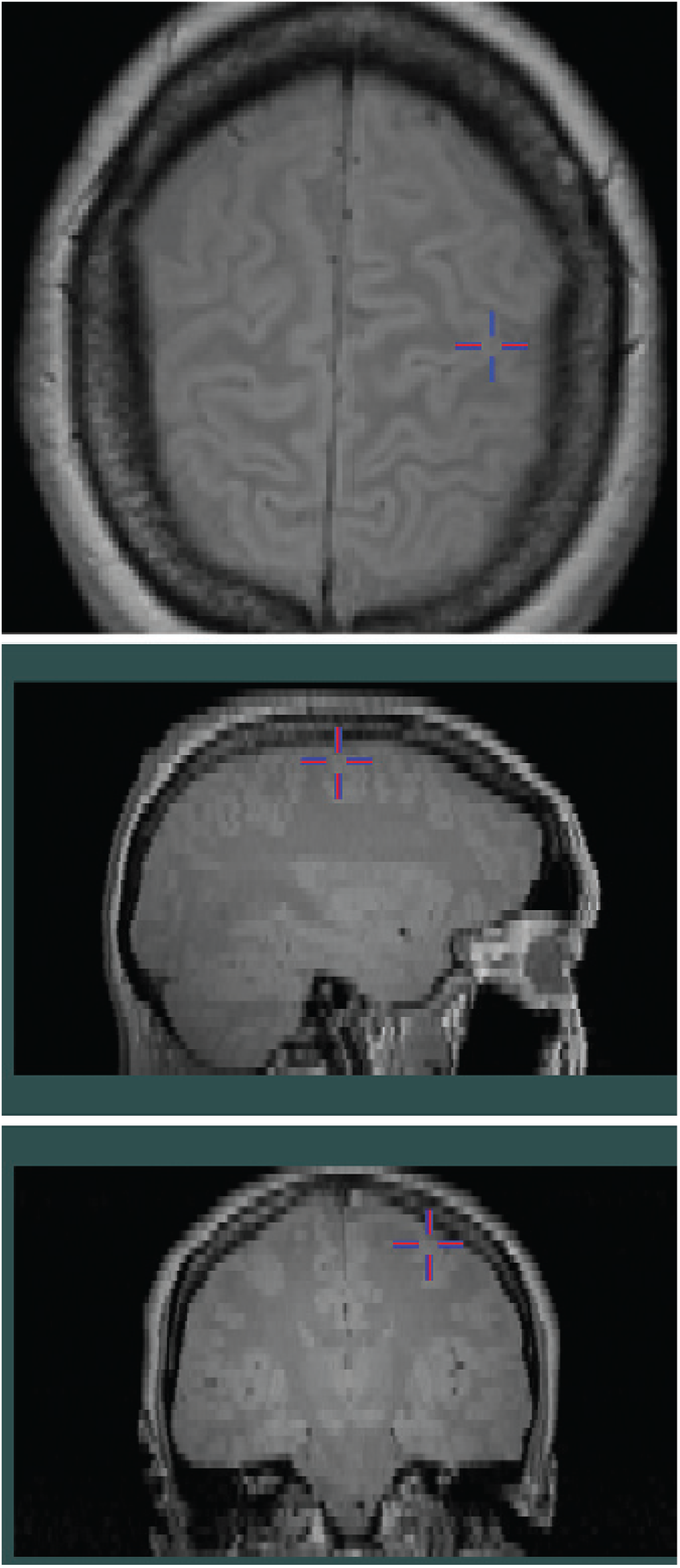
One of the 3 MRI datasets that were incorrectly classified as clean by the algorithm but had artifacts. A second visual inspection by an expert confirmed the artifacts. However, the expert noted that the artifacts were subtle compared to other datasets. The image on the right presents a proton density MRI contrast. The expert identified a very subtle motion artifact at the top.

The proposed DL algorithm benefited from the development over three stages, where we extended the size of the dataset the proportion of clean images, tuned the hyperparameters, and implemented MC dropout to generate probabilities and uncertainty measures. One of the biggest hurdles in this project was identifying a large dataset that was well-labeled with the ground truth: clean versus artifact. We overcame this challenge in stage I and into stage II where we extended the dataset to continue developing the algorithm. We added clean MRI volumes to increase the clean percentage from 15.9% to 50% by developing Clean Selector (Table A.10) to select scans constrained to the clean percentage by study. We used Clean Selector in stage III to extend the dataset to be 98% clean with over 30k MRI volumes, resembling other working large neuroimaging databases (Alfaro-Almagro et al., 2018), with some key differences:

- We used MRI volumes with multiple contrasts, while Alfaro-Almagro et al. (2018) used T1-weighted MRI only.
- Our dataset size was approximately 6 times larger, 34800/5816.
- Our data was acquired from various neuroimaging scanner manufacturers, while Alfaro-Almagro et al. (2018) used a single scanner manufacturer.

Despite this inherent variability in our data, the performance of the DL algorithm we developed outperformed all previous studies.

### 4.2. Learning rate and uncertainties

The LR is one of the most important hyperparameters to tune in DL. In stages II and III, we conducted two experiments (Fig. 3) to find a working LR range to implement cyclical LR. We customized the methods developed by Smith (2017) to develop our own decaying LR range. Our customized method provided sufficient freedom to the optimization process, with an initial wide ranging LR, and a final focusing period to fine-tune the network parameters, illustrated by the loss decrease in Figs. 7, 10, and 11.

In classification, the output generated by CNNs obtained by the softmax output layer, is commonly misinterpreted as model confidence. In stages II and III, we incorporated MC dropout to estimate probability and uncertainties. We used these metrics to filter MRI volumes with high uncertainty (i.e., send them to visual inspection) and select a probability-based threshold to optimize performance. Uncertainties proved useful in boosting the performance (Tables 6 and 7) and helpful in notifying the user of the prediction confidence. We estimated the number of inferences needed and found that increasing *N* resulted in decreased variance of the AUC estimates 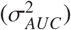 and decreased sum of squared errors (Fig. 14). At *N* = 10, the prediction already had small errors. To assess the performance, we used an even larger number of inferences, *N* > 50. Using uncertainties, we generated a wide-ranging set of results, highlighting the tradeoff between DFFRM and UDM (Fig. 22). The variety of experiments and uncertainties provide the users the flexibility to decide an appropriate operating point for their study. Where one user may wish to maximize the automation aspect of the quality control at the cost of higher UDM, another user may require low levels of UDM, and hire a team to visually inspect a large number of flagged MRI volumes.

### 4.3. The key in stage-wise development

Although they may appear trivial, stages I and II were necessary steppingstones to develop a robust high performing algorithm, that worked well not only with small balanced datasets but also with large and imbalanced datasets. Using the proof-of-concept dataset (Section 2.3.1) in stage I, we identified issues that need to be considered. We labeled the artifact MRI dataset and verified the clean MRI volumes were free from artifacts, giving us a well-labeled dataset and the confidence to continue developing the architecture for the proposed algorithm. After getting 100% accuracy in stage I, we could have decided that the problem was solved and reported our results. However, our goal was to create an algorithm that worked well with a large and imbalanced dataset.

To overcome the difference across domain labels in stage I (proof of concept), we applied ComBat to the dataset we used for this stage. As expected, the results in Fig. 16 present a lower performance relative to the performance obtained without applying ComBat (in Fig. 15). Regardless of the results, we conclude that using ComBat to harmonize the data removed the bias introduced by the study. In addition, the exceptional results obtained from the preliminary analysis before domain adaptation (Fig. 15) were biased due to having an exceptionally clean dataset from a single distinct trial. We would like to emphasize that we pursued stage I only for a proof of concept. The results of stage I do not influence the processing and results of stages II and III. However, for completeness, we now include the results we obtained while separately implementing stage I twice, without and with applying neuroCombat. Extending domain adaptation to stages II and III is not strictly necessary because to create the database we used for these stages we sampled over the same study distribution. That is, in stages II and III, we curated the dataset to remove the bias introduced in stage I by study, scanner, site, or scanning parameters. In these subsequent stages, there is no domain difference in either label or between cross-validation subsets. Both the artifact datasets and the clean datasets originated from the exact same distribution of scanners, pulse sequences, etc, Therefore, we believe that domain adaptation is not strictly required in stages II and III. Further, the very large datasets make it quite challenging to implement neuroCombat when the datasets need to be split into subsets of approximately 150 MRI scans. This is particularly true in practice where incoming datasets require additional pre-processing steps before detecting artifacts. That is, to implement neuroCombat, an additional harmonization step would be required at the beginning of the pipeline, resulting in another potential point of failure and increased chances for errors. Simplifying the automatic detection of artifacts by using native images directly has the potential to further improve the pipeline’s reliability.

Following stage I, we made naive attempts of using the realistic dataset (CP=98%), and obtained unimpressive results, with accuracy = 0.893 (specificity = 0.895 and sensitivity = 0.799; Fig. 6). Following these unsuccessful training sessions, we realized that using the realistic dataset to develop the algorithm was time consuming, and that a new approach was required. Instead, we created the balanced dataset (CP=50%) in stage II, allowing us to efficiently continue developing the methods presented in this paper.

In stage II, we developed Clean Selector to select clean images from the database, implemented cyclical LR to efficiently train the algorithm and incorporated MC dropout for probabilities and uncertainties. These additions allowed us to achieve high performance (Table 6) and extend to the realistic dataset. Our first attempt at transferring the learning from stage II to a realistic dataset by ramping the clean percentage did not yield high accuracies. That is, the linear increase in clean percentage resulted in a nonlinear increase in the dataset size and loss, as shown in Fig. 8. To control the dataset size and the amount of new data introduced to the algorithm we initiated data ramping experiments to compare methods for transferring the learning obtained in stage II.

The comparison experiments developed in stage III were designed to investigate how the final performance was affected by the initialization and the rate of ramping the dataset size. In Section 2.3.3, we formulated three experimental questions, addressed here. For Q1, amongst the two baseline experiments, when initialized with the stage II model, the algorithm achieved higher performance (Figs. 10 and 18). For Q2, the `S3_base_initialized` experiment generated comparable results to the data-ramping experiments (Figs. 18 and 22) suggesting that data-ramp provided a way for transfer learning, although comparable to initialization. For Q3, comparing amongst the data-ramping experiments revealed that using a lower ramp slope, `S3_dr_ER_100` performed best (Table 7, Fig. 18). To shed light on possible reasons for incorrect detection, we visually inspected specific cases of correct and incorrect classification. We present such cases in Section Appendix B of the Supplementary Material.

### 4.4. Contributions to Medical Imaging

We explored and exhausted several venues to culminate in the high-performing DL algorithm developed for detecting MRI artifacts in large imbalanced datasets. Our results in stage I gave us the confidence that we were on the right path, but rather misleading as our heedless jump to a realistic dataset proved not so straightforward. Making the decision to take a step back and create an intermediate balanced dataset proved to be extremely helpful. The smaller balanced dataset allowed us to explore several routes more efficiently, resulting in incorporating novel techniques outlined throughout the paper. Getting to the results in stage III was again not so simple, as we explored several routes to transfer the learning established in stage II. The comparison experiments were helpful in identifying that initialization and ramping the data slowly were key to transfer the learning from stage II. Other investigators may wish to use our approach. Of course, we expect that other creative techniques exist for transfer learning from a smaller dataset to a larger imbalanced dataset.

The comparison of the importance of the various aspects developed and explored in this project is summarized in Table 9. Minimal improvements were obtained with the LR schedule experiments. The techniques we developed and explored for handling class imbalance proved fruitful in obtaining improvements in performance across all metrics. Focal Loss reached performance levels obtained with transfer learning without data ramping; however, focal loss did not improve upon transfer learning with data ramping. This result emphasizes the relevance and importance of the methods we developed in this study. We note the discrepancy between the training and validation performances when using the focal loss, as apparent in Fig. 12. One explanation is that our model trained with the focal loss overfit the training dataset and thus did not learn features that generalize as well to unseen data. Regarding the performance obtained with MC dropout, the lowest metric in Table 9 is precision. However, in practice, this metric is related to the level of automation of our algorithm. A lower precision indicates a larger number of FP (clean MRI incorrectly classified to have artifacts), an inconvenience for database managers as it results in more MRI data being visually inspected. However, the critical number for database managers is FN (artifact MRI incorrectly classified to be clean) as these datasets cause the processing pipeline to generate errors that influence the conclusions. The performance for catching FN is well quantified with the recall metric. We demonstrate the capabilities of implementing aleatoric uncertainty to the proposed algorithm. Although using aleatoric uncertainty did not perform as well as using epistemic uncertainty, two possibilities could be explored further. One improvement could be to explore image processing techniques in addition to the axial flips that we implemented. As we discussed, these image processing techniques need to be implemented with care to not introduce artifacts to clean images. A second improvement could be to combine epistemic and aleatoric uncertainties to provide a new metric or threshold that can generate higher performance.

The results of this study have several contributions to the field of Medical Imaging and are summarized here. First, our proposed methods for handling an extreme class imbalance in detecting artifacts in a large MRI dataset outperform the use of focal loss. While both the focal loss and our approach operate at a high level of performance, the impact of the gap is amplified by the high incoming flow of MRI datasets that must be addressed in a timely manner. Using the methods presented in this paper, namely transfer learning, data-ramping, and MC dropout will provide high-quality tools needed for managing large imbalanced neuroimaging databases. Second, we evaluated variants of the novel data ramping method we developed to transfer the learning from a small balanced dataset to a large imbalanced dataset. We demonstrate that without this approach, there was a decrease in algorithm performance. Even by implementing focal cross-entropy, we were unable to improve the performance. Third, our study is the first to implement uncertainty measures in MRI artifact detection. Uncertainties form a powerful metric to assist in obtaining improved results and in practice give the users a confidence metric on the output generated by the algorithm. This confidence metric allows the user to make decisions on how to proceed. Fourth, the results we obtained supersede the performance of the automated methods that have been developed for artifact detection in large and imbalanced neuroimaging databases (Esteban et al., 2017; Pizarro et al., 2016; Alfaro-Almagro et al., 2018; Mortamet et al., 2009; Sujit et al., 2019). We conducted an in-depth comparison (Fig. 22) to the current state-of-the-art for quality detection, presented by Alfaro-Almagro et al. (2018) which was closest in dataset size and clean image percentage to the dataset we used. We expect that our DL approach supersedes their ML method because DL can extract informative features not explicitly defined for ML. Finally, the trend in neuroimaging has been to analyze larger neuroimaging datasets. We have demonstrated that despite obtaining great results with small datasets, it is necessary to make the transfer to large datasets carefully and thoughtfully.

## 5. Conclusion

We developed the first stochastic DL algorithm to generate automated high-performing MRI artifact detection as implemented on a large and imbalanced neuroimaging database. In stage I, we replicated the results of previous studies by obtaining 100% accuracy on a small dataset. The results from stage I confirmed that we could use DL for artifact detection. However, the dataset was not realistic in size and all clean scans were from a single study. In stages II and III, we developed Clean Selector to extend the size of the dataset and the fraction of clean MRI volumes (clean percentage), tuned the hyperparameters, and implemented Monte Carlo (MC) dropout to generate probabilities and uncertainty measures. Using a balanced dataset in stage II, we obtained 91.0% accuracy. By implementing uncertainty measures, we obtained 98.0% accuracy on 71.0% of the testing subset. In stage III, we devised a technique to transfer the learning obtained in stage II by ramping the data while asymptotically increasing the clean percentage. In stage III, we obtained 94.1% accuracy and by implementing uncertainty measures, we obtained 99.5% accuracy on 72.5% of the testing subset. We also framed our results as in (Alfaro-Almagro et al., 2018), using a dataset with similar size and fraction of clean MRI volumes. Compared to the results presented by Alfaro-Almagro et al. (2018), our algorithm provided improved alternatives: (i) similar artifact detection error with half the ‘manual work’ (visual inspection), or (ii) half the artifact detection error with the same amount of ‘manual work’ (visual inspection). The realistic dataset in stage III was larger than any previously reported work detecting artifact MRI. Our algorithm has been successfully implemented into the processing pipeline. The tool we developed has saved database management teams valuable resources, including hours spent by technicians for identifying artifacts in the MRI volume.

## Acknowledgments

This work was supported by the Mathematics of Information Technology and Complex Systems (Mitacs) Canada through the Mitacs Elevate grant. This research was undertaken thanks in part to funding from the Canada First Research Excellence Fund, awarded to McGill University for the Healthy Brains for Healthy Lives initiative. The work was supported by the Fonds de recherche du Qubec Sant.

We thank Professor Tal Arbel and her student Raghav Mehta for the helpful insight and discussions concerning the processing of a balanced dataset and implementing uncertainty metrics into deep learning.

## Software

The code for creating the network and the associated weights will be available at https://github.com/AS-Lab/Pizarro-et-al-2023-DL-detects-MRI-artifacts.

## Appendix A. Clean Selector

In subsection 2.1.2 we developed a process, entitled Clean Selector, illustrated in Table A.10 for the balanced 50% clean dataset, created in stage II. First, the artifact dataset distribution, **v**_*a*_, was used to compute the clean MRI volumes needed, 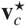, by setting 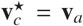, to maintain a study-wise balanced 50% clean dataset. Our plan was to take 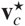 from the available clean MRI volumes, 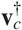. However, as quantified in Table A.10, for studies *s*\* = [3, 6, 16, 18, 20] no clean MRI volume was available, i.e.,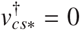. We therefore computed the remainder, 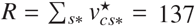, as the sum of the clean MRI volume needed to maintain the overall dataset at cp=50%. We then used the artifact dataset distribution, **v**_*a*_ to redistribute *R* as **v**_*R*_, setting *v*_*Rs*\*_ = 0. Finally, we computed the clean MRI taken, **v**_*c*_, as follows:

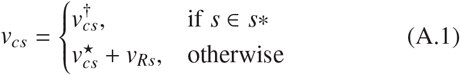

with final study-wise clean percent, *cp*_*s*_ = *v*_*cs*_/(*v*_*as*_ + *v*_*cs*_). This two-step process allowed us to efficiently identify the number of MRI volumes needed to maintain a study-wise 50% clean dataset, based on the available data.

In stage III, we used Clean Selector to determine MRI volumes taken by study to extend the dataset to 98% clean, a proportion reflecting reality. First, the artifact dataset distribution, **v**_*a*_, was used to compute the number of clean MRI volumes needed, 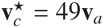, to maintain a study-wise 98% clean dataset. However, the available clean MRI volumes, 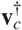, had the same constraint as in stage II, i.e., for *s*\* = [3, 6, 16, 18, 20], 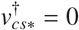. In addition, 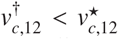, i.e., for study *s* = 12, the clean MRI available was smaller than the clean MRI needed. We therefore computed the remainder, 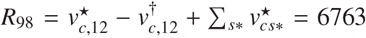 as the sum of the clean MRI volumes needed to maintain the overall dataset at cp=98%. We then used the artifact dataset distribution, **v**_*a*_ to redistribute *R*_98_ as **v**_*R*_, setting *v*_*Rs*\*_ = *v*_*R*,12_ = 0 and computed the clean MRI taken **v**_*c*_, using Eq. A.1.

## Appendix B. Secondary visual-inspection

To shed light on possible reasons for incorrect detection, we visually inspected specific cases of correct and incorrect classification. We used the results from Table 7, row σ^2^ < 0.01; we present these results here as a confusion matrix (Fig. B.24) as context for the discussion. With an MRI expert, we visually inspected the five images with artifacts that the algorithm classified as clean images (false negative, FN=5, top right of confusion matrix). We note that FNs are the most problematic cases because these MRI volumes with artifacts would be incorrectly classified as clean, and they will be sent for processing in the pipeline. In other words, the analysis and conclusions will be based not only on clean images but also on images with artifacts. These were the cases we aimed to minimize.

This secondary visual inspection by the expert revealed that two MRI datasets out of the five that were previously labeled as artifacts were indeed clean and acceptable for analysis. The expert did not agree with the initial human rater’s label that the MRI images showed movement artifacts. The two MRI contrasts were magnetic transfer ON and proton density (Pizarro et al., 2019). We include an image for one of these two MRI volumes in Fig. B.25. The remaining three out of five MRI datasets had artifacts and were incorrectly classified by the algorithm, although the expert noted that the artifact was subtle compared to other datasets (Fig. B.26). Regardless, this would cause a problem in the processing pipeline. We include in Fig. B.26 one such MRI volume that was misclassified as clean.

In addition, we visually inspected the 25 clean images that were classified by the algorithm as images with artifacts (false positive, FP = 25, bottom left of confusion matrix). We note that FP cases are less problematic than FN cases. In practice, these clean MRIs would be incorrectly labeled as artifacts and get sent for visual inspection, at which point the MRI expert would relabel them as clean. We categorized the results of this visual inspection as follows:

- 11 clean cases that were labeled clean were confirmed to be clean and free from artifacts in the second inspection by the expert. These images were incorrectly classified by the algorithm as showing artifacts.
- 8 cases that were previously labeled as clean had artifacts, as confirmed by the expert in the second visual inspection. The algorithm correctly identified them as images with artifacts.
- 6 cases were labeled as clean but contained a relatively large number of abnormal lesions. These lesions confused the algorithm and therefore the images were classified as having artifacts.

This secondary in-depth visual inspection revealed some potential venues to improve the classifier in the future. In our current study, we selected MRI volumes with an artifact from those with prominent artifacts. Given the capability of generating uncertainty and probability values with MC dropout, future directions may include training the algorithm with datasets that have subtle artifacts. These images with subtle artifacts would be labeled with a value in the range of [0, 1]. After training with these images that have subtle artifacts, the algorithm should be capable of generating probability values of having an artifact, including relatively low probabilities associated with subtle artifacts. This would then reduce the errors for the images that have subtle artifacts. In addition, it is possible to provide more examples of images with lesions labeled as clean MRI by incorporating data from publicly available databases, to avoid classification of such images as images with artifacts.

## Footnotes

1 Abbreviations used throughout the text are summarized in Table 1

## References

Aleef, T.A., Spadinger, I.T., Peacock, M.D., Salcudean, S.E., Mahdavi, S.S., 2021. Rapid treatment planning for low-dose-rate prostate brachytherapy with tp-gan, in: International Conference on Medical Image Computing and Computer-Assisted Intervention, Springer. pp. 581–590.

Alfaro-Almagro, F., Jenkinson, M., Bangerter, N.K., Andersson, J.L.R., Griffanti, L., Douaud, G., Sotiropoulos, S.N., Jbabdi, S., Hernandez-Fernandez, M., Vallee, E., Vidaurre, D., Webster, M., McCarthy, P., Rorden, C., Daducci, A., Alexander, D.C., Zhang, H., Dragonu, I., Matthews, P.M., Miller, K.L., Smith, S.M., 2018. Image processing and quality control for the first 10,000 brain imaging datasets from UK biobank. Neuroimage 166, 400–424.

Chen, J., Wan, Z., Zhang, J., Li, W., Chen, Y., Li, Y., Duan, Y., 2021. Medical image segmentation and reconstruction of prostate tumor based on 3d alexnet. Computer methods and programs in biomedicine 200, 105878.

Chollet, F., 2017. Deep Learning with Python. Manning Publications.

Das, S., Zijdenbos, A.P., Harlap, J., Vins, D., Evans, A.C., 2012. LORIS: a web-based data management system for multi-center studies. Frontiers in Neuroinformatics 5.

Di Martino, A., Yan, C.G., Li, Q., Denio, E., Castellanos, F.X., Alaerts, K., Anderson, J.S., Assaf, M., Bookheimer, S.Y., Dapretto, M., Deen, B., Delmonte, S., Dinstein, I., Ertl-Wagner, B., Fair, D.A., Gallagher, L., Kennedy, D.P., Keown, C.L., Keysers, C., Lainhart, J.E., Lord, C., Luna, B., Menon, V., Minshew, N.J., Monk, C.S., Mueller, S., Müller, R.A., Nebel, M.B., Nigg, J.T., O’Hearn, K., Pelphrey, K.A., Peltier, S.J., Rudie, J.D., Sunaert, S., Thioux, M., Tyszka, J.M., Uddin, L.Q., Verhoeven, J.S., Wenderoth, N., Wiggins, J.L., Mostofsky, S.H., Milham, M.P., 2014. The autism brain imaging data exchange: towards a large-scale evaluation of the intrinsic brain architecture in autism. Mol. Psychiatry 19, 659–667.

Ducharme, S., Albaugh, M.D., Nguyen, T.V., Hudziak, J.J., Mateos-Pérez, J.M., Labbe, A., Evans, A.C., Karama, S., 2016. Trajectories of cortical thickness maturation in normal brain development — the importance of quality control procedures. NeuroImage 125, 267–279.

Esteban, O., Birman, D., Schaer, M., Koyejo, O.O., Poldrack, R.A., Gorgolewski, K.J., 2017. MRIQC: Advancing the automatic prediction of image quality in MRI from unseen sites. PLoS One 12, e0184661.

Fantini, I., Rittner, L., Yasuda, C., Lotufo, R., 2018. Automatic detection of motion artifacts on MRI using deep CNN, in: 2018 International Workshop on Pattern Recognition in Neuroimaging (PRNI), pp. 1–4.

Fernandez, V.G., Asarnow, R., Narr, K.L., Subotnik, K.L., Kuppinger, H., Fogelson, D., Nuechterlein, K.H., 2018. Temporal lobe thickness and verbal memory in first-degree relatives of individuals with schizophrenia. Schizophr. Res. 199, 221–225.

Fortin, J.P., Cullen, N., Sheline, Y.I., Taylor, W.D., Aselcioglu, I., Cook, P.A., Adams, P., Cooper, C., Fava, M., McGrath, P.J., McInnis, M., Phillips, M.L., Trivedi, M.H., Weissman, M.M., Shinohara, R.T., 2018. Harmonization of cortical thickness measurements across scanners and sites. NeuroImage 167, 104–120. URL: https://www.sciencedirect.com/science/article/pii/S105381191730931X, doi:https://doi.org/10.1016/j.neuroimage.2017.11.024.

Gal, Y., Ghahramani, Z., 2016. Dropout as a bayesian approximation: Representing model uncertainty in deep learning, in: international conference on machine learning, pp. 1050–1059.

Gardner, E.A., Ellis, J.H., Hyde, R.J., Aisen, A.M., Quint, D.J., Carson, P.L., 1995. Detection of degradation of magnetic resonance (MR) images: comparison of an automated MR image-quality analysis system with trained human observers. Acad. Radiol. 2, 277–281.

Herzog, L., Murina, E., Dürr, O., Wegener, S., Sick, B., 2020. Integrating uncertainty in deep neural networks for MRI based stroke analysis. Med. Image Anal. 65, 101790.

Japkowicz, N., Stephen, S., 2002. The class imbalance problem: A systematic study1. Intelligent Data Analysis 6, 429–449.

Jubault, T., Gagnon, J.F., Karama, S., Ptito, A., Lafontaine, A.L., Evans, A.C., Monchi, O., 2011. Patterns of cortical thickness and surface area in early parkinson’s disease. Neuroimage 55, 462–467.

Kendall, A., Gal, Y., 2017. What uncertainties do we need in bayesian deep learning for computer vision? Advances in neural information processing systems 30.

Knopman, D.S., Lundt, E.S., Therneau, T.M., Vemuri, P., Lowe, V.J., Kantarci, K., Gunter, J.L., Senjem, M.L., Mielke, M.M., Machulda, M.M., Roberts, R.O., Boeve, B.F., Jones, D.T., Petersen, R.C., Jack, Jr, C.R., 2018. Joint associations of β-amyloidosis and cortical thickness with cognition. Neurobiol. Aging 65, 121–131.

Küstner, T., Liebgott, A., Mauch, L., Martirosian, P., Bamberg, F., Nikolaou, K., Yang, B., Schick, F., Gatidis, S., 2018. Automated reference-free detection of motion artifacts in magnetic resonance images. MAGMA 31, 243–256.

Lin, T.Y., Goyal, P., Girshick, R., He, K., Dollár, P., 2017. Focal loss for dense object detection, in: Proceedings of the IEEE international conference on computer vision, pp. 2980–2988.

Liu, Z., Mao, H., Wu, C.Y., Feichtenhofer, C., Darrell, T., Xie, S., 2022. A convnet for the 2020s, in: Proceedings of the IEEE/CVF Conference on Computer Vision and Pattern Recognition, pp. 11976–11986.

Marcus, D.S., Harms, M.P., Snyder, A.Z., Jenkinson, M., Wilson, J.A., Glasser, M.F., Barch, D.M., Archie, K.A., Burgess, G.C., Ramaratnam, M., Hodge, M., Horton, W., Herrick, R., Olsen, T., McKay, M., House, M., Hileman, M., Reid, E., Harwell, J., Coalson, T., Schindler, J., Elam, J.S., Curtiss, S.W., Van Essen, D.C., WU-Minn HCP Consortium, 2013. Human connectome project informatics: quality control, database services, and data visualization. Neuroimage 80, 202–219.

de Mathelin, A., Deheeger, F., Richard, G., Mougeot, M., Vayatis, N., 2021. Adapt: Awesome domain adaptation python toolbox. arXiv preprint arXiv:2107.03049.

Menon, S., Chapman, D., Nguyen, P., Yesha, Y., Morris, M., Saboury, B., 2020. Deep expectation-maximization for semi-supervised lung cancer screening. arXiv preprint arXiv:2010.01173.

Mortamet, B., Bernstein, M.A., Jack, C.R., Gunter, J.L., Ward, C., Britson, P.J., Meuli, R., Thiran, J.P., Krueger, G., Alzheimer’s Disease Neuroimaging Initiative, 2009. Automatic quality assessment in structural brain magnetic resonance imaging. Magnetic Resonance in Medicine 62, 365–372.

Munappy, A., Bosch, J., Olsson, H.H., Arpteg, A., Brinne, B., 2019. Data management challenges for deep learning, in: 2019 45th Euromicro Conference on Software Engineering and Advanced Applications (SEAA), pp. 140–147.

Nair, T., Precup, D., Arnold, D.L., Arbel, T., 2020. Exploring uncertainty measures in deep networks for multiple sclerosis lesion detection and segmentation. Med. Image Anal. 59, 101557.

Najafabadi, M.M., Villanustre, F., Khoshgoftaar, T.M., Seliya, N., Wald, R., Muharemagic, E., 2015. Deep learning applications and challenges in big data analytics. Journal of Big Data 2.

Oksuz, I., 2020. Brain MRI artefact detection and correction using convolutional neural networks. Comput. Methods Programs Biomed. 199, 105909.

Pizarro, R., Assemlal, H.E., De Nigris, D., Elliott, C., Antel, S., Arnold, D., Shmuel, A., 2019. Using deep learning algorithms to automatically identify the brain MRI contrast: Implications for managing large databases. Neuroinformatics 17, 115–130.

Pizarro, R.A., Cheng, X., Barnett, A., Lemaitre, H., Verchinski, B.A., Goldman, A.L., Xiao, E., Luo, Q., Berman, K.F., Callicott, J.H., Weinberger, D.R., Mattay, V.S., 2016. Automated quality assessment of structural magnetic resonance brain images based on a supervised machine learning algorithm. Front. Neuroinform. 10, 52.

Rani, S., Ghai, D., Kumar, S., Kantipudi, M., Alharbi, A.H., Ullah, M.A., 2022. Efficient 3d alexnet architecture for object recognition using syntactic patterns from medical images. Computational Intelligence and Neuroscience 2022.

Reuter, M., Tisdall, M.D., Qureshi, A., Buckner, R.L., van der Kouwe, A.J.W., Fischl, B., 2015. Head motion during MRI acquisition reduces gray matter volume and thickness estimates. Neuroimage 107, 107–115.

Shamsi, A., Asgharnezhad, H., Abdar, M., Tajally, A., Khosravi, A., Nahavandi, S., Leung, H., 2021. Improving mc-dropout uncertainty estimates with calibration error-based optimization. arXiv preprint arXiv:2110.03260

Smith, L.N., 2017. Cyclical learning rates for training neural networks, in: 2017 IEEE Winter Conference on Applications of Computer Vision (WACV), pp. 464–472.

Sujit, S.J., Coronado, I., Kamali, A., Narayana, P.A., Gabr, R.E., 2019. Automated image quality evaluation of structural brain MRI using an ensemble of deep learning networks. J. Magn. Reson. Imaging 50, 1260–1267.

Tanno, R., Worrall, D.E., Kaden, E., Ghosh, A., Grussu, F., Bizzi, A., Sotiropoulos, S.N., Criminisi, A., Alexander, D.C., 2021. Uncertainty modelling in deep learning for safer neuroimage enhancement: Demonstration in diffusion MRI. Neuroimage 225, 117366.

Yau, Y., Zeighami, Y., Baker, T.E., Larcher, K., Vainik, U., Dadar, M., Fonov, V.S., Hagmann, P., Griffa, A., Mišić, B., Collins, D.L., Dagher, A., 2018. Network connectivity determines cortical thinning in early parkinson’s disease progression. Nat. Commun. 9, 12.

Yu, Y., Li, M., Liu, L., Li, Y., Wang, J., 2019. Clinical big data and deep learning: Applications, challenges, and future outlooks. Big Data Mining and Analytics 2, 288–305.

Zhao, G., Liu, F., Oler, J.A., Meyerand, M.E., Kalin, N.H., Birn, R.M., 2018. Bayesian convolutional neural network based MRI brain extraction on nonhuman primates. NeuroImage 175, 32–44.

